# Microbial Responses to Biochar Soil Amendment and Influential Factors: A Three-level Meta-analysis

**DOI:** 10.1101/2023.06.02.543269

**Authors:** Patricia Kerner, Ethan Struhs, Amin Mirkouei, Ken Aho, Kathleen A Lohse, Robert S Dungan, Yaqi You

## Abstract

Biochar is a multifunctional soil conditioner capable of enhancing soil health and crop production while reducing greenhouse gas emissions. Understanding how soil microbes respond to biochar amendment is a vital step towards precision biochar application. Here, we synthesized 3899 observations of 24 microbial responses from 61 primary studies, applied a three-level mixed-effects model to estimate biochar effects, and evaluated the importance of biochar characteristics (feedstock, pyrolysis temperature), soil properties (pH, C:N, cation exchange capacity, bulk or rhizosphere), and treatment protocols (application rate, fertilization, duration, field or laboratory). Biochar significantly boosts microbial abundance (microbial biomass carbon > CFU), nitrite reductase gene (*nirS*), the activity of C- and N-cycling enzymes (dehydrogenase > cellulase > urease > invertase), and potential nitrification rate. Biochar characteristics, soil properties, and treatment protocols strongly determine the direction and extent of microbial response changes. Feedstock, pyrolysis temperature, application rate, and soil pH are important predictors most frequently included in the final models. Our study highlights the promise of purpose-driven biochar production and application such that biochar production parameters can be tuned to elicit the desired microbial responses and application protocols could be optimized to invoke multiple benefits. It also underlines current knowledge gaps and future research needs.

**Synopsis:** Meta-analysis reveals overall effect sizes of soil microbial responses to biochar amendment and the most influential factors, highlighting the potential of purpose-driven precision biochar towards sustainable agriculture.

## 1. Introduction

Soil is a vital ecosystem that sustains food security and other development goals.^1^ During the past decade, biochar has received increasing attention due to its promise as a low-cost, multifunctional soil conditioner (Figure S1).^2^ Numerous studies have reported that biochar soil amendment is capable of preserving or improving soil quality, promoting crop production, decreasing nutrient leaching, and reducing greenhouse gas (GHG) emissions from agricultural soils.^3–5^ Yet the mechanisms underlying biochar’s beneficial effects on the soil-plant system are not fully understood, which hinders the realization of the full potential of biochar in sustainable soil management.

Soil contains a vast diversity of microorganisms that together mediate soil functions and directly contribute to plant fitness in a changing environment.^6,7^ Biochar amendment can alter the indigenous soil microbiome, which could in turn drive shifts in soil functionality. Indeed, studies around the world have identified significant alterations in soil microbial biomass, diversity, community composition, and enzyme activity following biochar addition (e.g., refs^8–11^). Some, however, found only minor changes in these perspectives (e.g., refs ^12–14^).

The large variations and sometimes contrasting outcomes across studies could be attributed to heterogeneity in soil characteristics, biochar properties, or treatment protocols. First, soil is a complex matrix varying in abiotic properties (e.g., pH, cation exchange capacity (CEC)) and biotic components (e.g., the indigenous soil microbiome). Second, biochar can be derived from a wide range of organic materials (i.e., feedstocks) under oxygen-limiting conditions. Biochar physicochemical properties, such as porosity, aromaticity, and surface functional groups, vary considerably depending on the feedstock and conversion process.^15,16^ Third, the actual treatment often differs substantially in terms of treatment frequency, rate, and duration, among other conditions.

A systematic review of the current literature would allow us to synthesize knowledge of soil microbiome responses to biochar amendment, unveil key factors that can influence biochar effects on the soil microbiome, and identify significant knowledge gap in the field. Specifically, meta-analysis quantitatively analyzes the results from multiple primary studies, allowing more robust conclusions to be drawn and new insights to be gained.^17^ Meta-analysis has been employed to investigate biochar effects on soil quality,^18^ crop production,^4^ GHG emission,^19,20^ and more recently also some microbial parameters.^11,21–23^ However, influential factors of microbial responses are not explored.

Here we utilized a three-level meta-analysis framework to quantitatively synthesize the most recent global findings of biochar impacts on the soil microbiome, examined the influence of biochar characteristics, soil properties, and treatment protocols on soil microbial responses, and ranked potential drivers of microbial responses at the molecular, population and community level. This framework has a hierarchical structure of an extended mixed-effects model, allowing dependent observations to be retained while heterogeneity at different levels being quantitatively assessed.^24,25^ Our results provide novel insights into biochar’s benefits to the soil-plant system and underscore the promise of purpose-driven biochar production and application towards sustainable agriculture.

## 2. Materials and Methods

### 2.1. Systematic Literature Review

A systematic literature search was performed using the keyword “biochar” paired with soil microbial response terms (listed in Table S1) in the Web of Science to identify relevant peer-reviewed articles. Publications were limited by date (between January 2018 and April 2020) and manuscript type (original research). This resulted in an initial collection of 3511 publications, from which 1777 duplicates were removed. Publications were further screened according to the following inclusion criteria: 1) experimental design allowed pairwise comparison between biochar treatment and no-biochar control, 2) if fertilizer was applied, both biochar-only and no-fertilizer controls were available to isolate the effects of biochar, 3) soil used was not contaminated with metal or mine waste, 4) biochar was produced by pyrolyzing organic materials under dry conditions and was not preincubated, 5) microbial inoculants were absent, and 6) the study reported sample sizes, means and standard errors or the information was given by the author through personal communication. This resulted in 61 studies (see SI references), including 14 field studies, and 3899 pairwise treatment-control comparisons. Figure 1 shows a PRISM (Preferred Reporting Items for Systematic Reviews and Meta-Analyses) diagram summarizing the screening procedure.^26^ More details are in the Supporting Information (SI).

**Figure 1.**
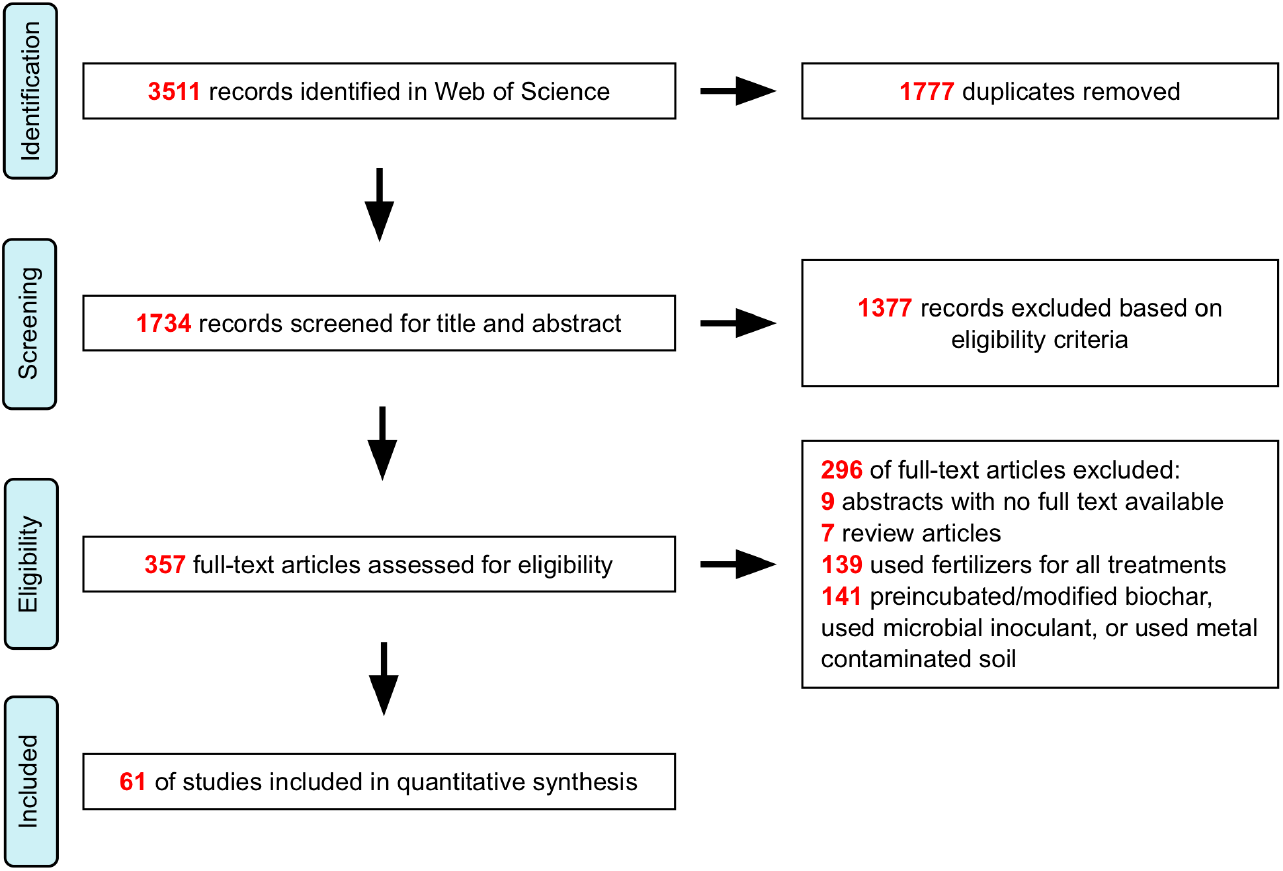
PRISMA diagram summarizing the number of studies screened for and retained through this meta-analysis.

### 2.2. Data Extraction

We examined the 61 studies carefully to extract data including experimental designs, biochar production conditions and characteristics, soil properties, soil biogeochemistry, soil (including rhizosphere) microbial responses, and plant responses when available. Data were collected directly from original tables where possible; data only presented in figures were extracted using WebPlotDigitizer (v4.2), or from authors directly. A variety of response variables were obtained (Table S2) and those with >=20 observations were included in the meta-analysis.

Similar to others,^19,27^ we initially assigned predictor variables (moderators) to categorical groups to facilitate cross-study comparisons, considering the distribution of categorical levels in the entire database (Table S3). Biochar feedstocks were grouped into the following categories: (1) manure (poultry, pig or cattle manure), (2) sludge (water treatment plant sewage sludge), (3) wood (hardwoods such as pine, oak, beech, fir and bamboo, and wood mixtures), (4) agricultural biomass (residues from rice, wheat, corn, sugarcane and legumes), and (5) lignocellulose (nut shells, fruit peels, weeds and tree leaves). If soil texture was not reported in a study, the harmonized world soil database^19,27,28^ was used to extract soil texture information according to the reported coordinates. When soil organic matter (SOM) rather than soil organic carbon (SOC) was reported, SOC was estimated as 58% of SOM.^29^ Biochar application rates reported as metric tons/hectare were converted to mass percentage using the reported soil bulk density.

### 2.3. Three-level Meta-analytical Model

We followed the general methodology in Koricheva et al. (2013),^30^ from data compiling to statistical modeling, to conduct a meta-analysis on the extracted data. The effect sizes of biochar treatments on soil microbial responses, as well as soil or plant responses, were measured using the log response ratio (*LRR*):^31^

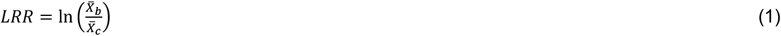

Where 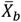 is the mean of a response variable under biochar treatment and 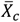 is the mean of the response variable under the control.

The estimate of study variance 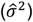 was calculated using the reported variance of the mean of a response variable:

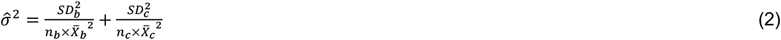

Where *SD*^2^ indicates the reported standard deviation of the mean of the response variable and *n* is the sample size or reported number of replicates. We used a three-level random-effects model for effect sizes, assuming random effects at different levels and independent sampling error:^32^

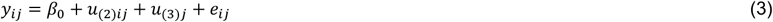

Where *y*_*ij*,_is the *i*th effect size in the *j*th study; *β*_0_is average population effect; *Var*(*e*_*ij*_),*= v*_*ij*,_ with *e*_*ij*,_ being the sampling error of the *i*th effect size in the *j*th study (level 1); 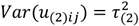 is the within-study heterogeneity (level 2); 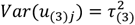 is the between-study heterogeneity (level 3). We assumed that the marginal errors and random effects were normally distributed with a mean of 0, and were independent. We used the statistic *I*^*2*^ to estimate the proportion of variation in effect sizes explained by level-2 or level-3 variance, with total heterogeneity being the sum of both,^33^ and Cochran’s Q to test the significance of heterogeneity (also see the SI).^34^ Model parameters were estimated using restricted maximum likelihood (REML).^35^ Sources of high *I*^*2*^ were explored through extending Eq. 3 to a three-level mixed-effects model with individual moderators as a covariate:^36^

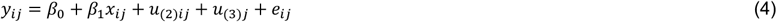

Where *β*_1_ is a moderator regression coefficient; *x* is a moderator covariate at either level 2 (*x*_*ij*,_) or level 3 (*x*_,*j*_; same for all effect sizes in the *j*th study); conditional mean *E*(*y*_*ij*,_|*x*_*ij*,_) *= β* + *β*_1_*x*_*ij*,_; conditional variance 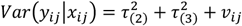 conditional covariances 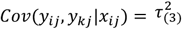 (i.e., the same covariance is shared by effect sizes in the *j*th study); *Cov*(*y*_*ij*,_, *y*_mn,_|*x*_*ij*_), *=* 0 (i.e., independent effect sizes are assumed for different studies); 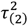 and 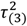 are the level-2 and level-3 residual heterogeneity after controlling for the covariate. Nine moderators were considered: biochar characteristics (feedstock, pyrolysis temperature), soil properties (pH, C:N, CEC, bulk or rhizosphere), treatment protocols (biochar application rate, fertilization, experiment duration, field or laboratory). After analyzing individual moderators, we extended the three-level mixed-effects model to contain multiple moderators. Here soil pH, soil C:N, experiment duration and biochar application rate were treated as continuous predictors, while others were categorical. Soil type, fertilization and experiment type were excluded due to the low number of observations and/or overlapping with other moderators. Meta-regression models considering only main effects (i.e., without interactions) were fitted based on log-likelihoods. Multicollinearity of candidate predictors was evaluated by using variance inflation factor (VIF). Non-correlated predictors were included in automated model selection for each response variable. The most parsimonious model was selected using the corrected Akaike information criterion (AICc), and model-averaged moderator importance was also calculated. Maximum likelihood estimation instead of REML was used to allow the use of AIC to compare for models with different fixed effects.^37^ The percentage of heterogeneity explained by the inclusion of one or more moderators was estimated using a pseudo-*R*^2^ statistics:^38^

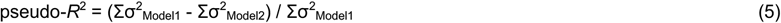

Where Σσ^2^_Model1_ is the sum of residual variances of the random intercept components in the overall model (no moderator) and Σσ^2^_Model2_ is the sum of residual variances of the random intercept components in a model with one or more moderators for a given response variable. If the pseudo-*R*^*2*^ calculation generated negative values, they were truncated to zero. Additional modeling details including multimodel inference are in the SI.

### 2.4. Publication Bias and Missing Data

Techniques for handling missing data in conventional meta-analysis have not been comprehensively evaluated for multi-level meta-analysis.^36^ Here we utilized multiple techniques, before and after model selection, to better assess and handle missing data due to publication bias (details in the SI).

### 2.5. Computational Implementation

All analysis was performed in R (v3.6.1) with package ‘metafor’ for meta-analysis, ‘corrplot’ and ‘performance’ for collinearity examination, ‘glmulti’ for automated stepwise model selection, and ‘ggplot2’ for data visualization.^38–42^

## 3. Results and Discussion

### 3.1. Global Estimate of Microbial Responses

We obtained twenty-four response variables with ≥20 observations after examining 61 primary studies and 3899 pairwise treatment-control comparisons (Table S4). These included microbial abundance (bacterial colony forming units (CFU), bacterial phospholipid fatty acid (PLFA), fungal PLFA, microbial biomass carbon (MBC), microbial biomass nitrogen (MBN), diversity (ACE, Chao1, Shannon, Simpson), gene abundance (archaeal *amoA*, bacterial *amoA, narG, nirS, nosZ*), enzyme activity (acid phosphatase, alkaline phosphatase, β-glucosidase, cellulase, dehydrogenase, invertase, urease), and process (cumulative CO_2_, cumulative N_2_O, potential nitrification rate). The genes, enzymes, and processes are involved in C, N, and P cycling. Except for CFU and ACE, our three-level random-effects model explained 77.7%-99.9% of the total heterogeneity in these variables (in the SI).

Biochar soil amendment increased 21 out of the 24 variables (Figure 2; Table S5). Increases were significant for two abundance variables CFU (+1.74%, n = 32) and MBC (+26.5%, n = 202), the prevalence of nitrite reductase gene *nirS* (+8.2%, n = 28), the activity of cellulase (+55.6%, n = 40), dehydrogenase (+84.1%, n = 128), invertase (+21.2%, n = 31) and urease (+39.4%, n = 74), and potential nitrification rate (+40.8%, n = 33). Biochar insignificantly decreased 3 variables: MBN (−4.0%, n = 116), ammonium-oxidizing archaea (AOA) (−17.4%, n = 28), and cumulative N_2_O (−12.7%, n = 71). Ammonium-oxidizing bacteria (AOB) had the largest response variance, ranging from -54.8% to +351.8% (n = 61, 5 studies), which was not due to within- or between study variation alone (49.4% and 50.6% of total heterogeneity, respectively). Overall, our global estimates highlight that biochar can modulate the soil microbiome, resulting in a wide range of alterations at the molecular, population and community level.

**Figure 2.**
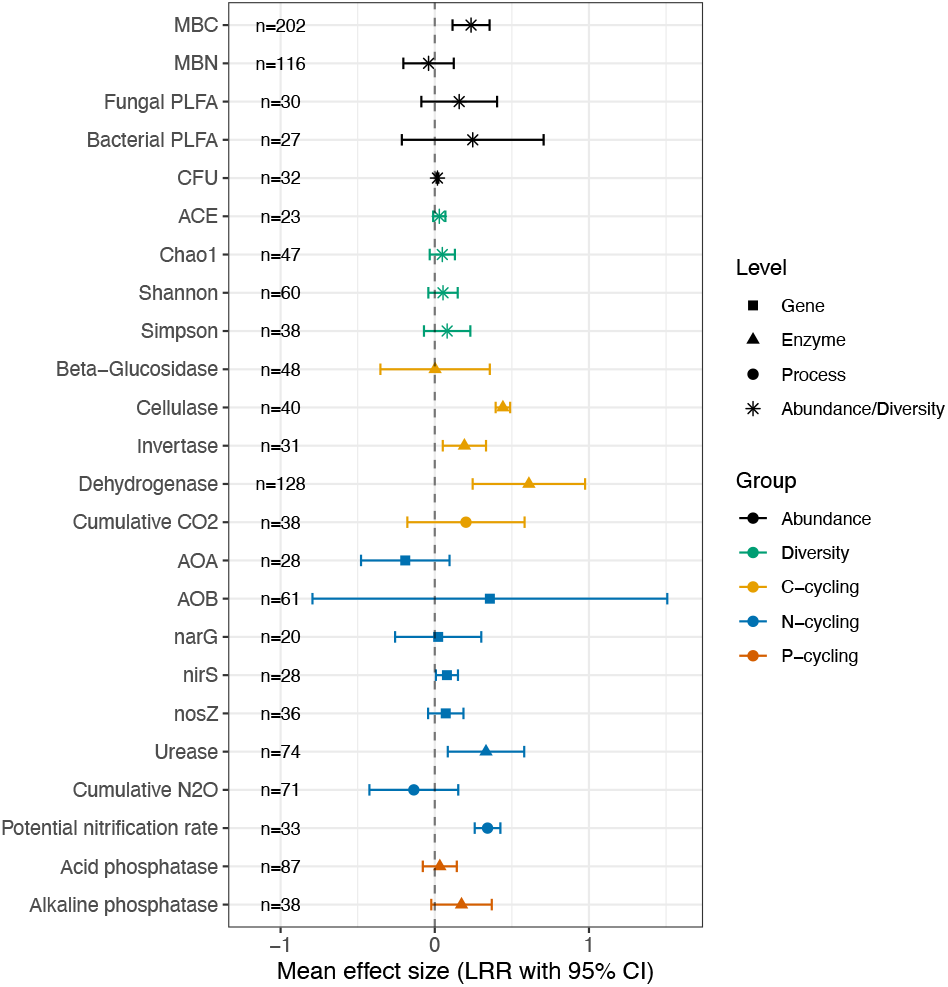
Mean effects of biochar on soil microbiome responses (weighted LRR ± 95% confidence interval). Effect sizes where 95% confidence intervals do not overlap with zero are significant. AOA, ammonium-oxidizing archaea; AOB, ammonium-oxidizing bacteria; CFU, colony forming unit; MBC, microbial biomass carbon; MBN, microbial biomass nitrogen; PLFA, phospholipid fatty acid. AOB and AOA were commonly measured with quantitative PCR targeting the ammonia monooxygenase gene *amoA*.

### 3.2. Biochar Effects on Soil C, N, P Cycling

Soil microbiome alterations could explain biochar’s conditioner functionality. Biochar significantly stimulated the activity of three C-cycling enzymes, cellulase, dehydrogenase, and invertase (Figure 2; Table S5). Cellulase decomposes cellulose and related polysaccharides, responsible for the turnover of plant biomass in the biosphere. Three major types are produced by certain bacteria and fungi: 1,4-β-cellobiosidase (EC 3.2.1.91/176), endo-1,4-β-D-glucanase (EC 3.2.1.4), and β-glucosidase (EC 3.1.2.21).^43^ Biochar enhanced the activity of cellulase in general, but had no significant effect on the activity of β-glucosidase that completes the final step of cellulose hydrolysis. Strongest enhancement was observed in intracellular dehydrogenase (EC 1.1.1.) that catalyzes soil organic matter oxidation. Dehydrogenase activity is proportional to microbial biomass.^44,45^ Biochar’s strong stimulation of dehydrogenase activity is likely attributed to improved soil aeration, water availability, pH, and nutrient availability that are known to significantly influence dehydrogenase activity.^16,44,46^ These results are partially consistent with two other meta-analyses that focused specifically on enzyme activities, which found that biochar had insignificant or negative effect on 1,4-β-cellobiosidase, insignificant effect on β-glucosidase, and positive effect on dehydrogenase.^9,11^ It should be noted that our analysis included all the three types of cellulases, representing a more comprehensive overview. Fewer studies have investigated invertase (EC 3.2.1.26), which is widely distributed in microorganisms, both intracellular and extracellular, and attacks β-D-fructofuranoside in oligosaccharides present in vegetables.^47^ Individual studies have reported both increased and decreased invertase activity, depending on soil type, biochar dose, and experimental design.^48,49^ Our results based on data from 7 studies suggested generally enhanced invertase activity. Higher activity of C-cycling enzymes may result in faster C turnover and our estimate showed a 22.4% increase in cumulative CO_2_, while another report suggested 19% increase.^20^ These results are further supported by a recent study showing biochar-induced bacterial mineralization of soil recalcitrant components.^50^

Among the N-cycling responses, biochar significantly stimulated *nirS* gene, the activity of urease (EC 3.5.1.5) that hydrolyzes urea to ammonia and CO_2_, and potential nitrification rate. Meanwhile, biochar reduced AOA and cumulative N_2_O although insignificantly. N cycle is driven by complex networks of metabolically versatile microorganisms.^51,52^ Biochar has been shown to increase urease activity, AOA abundance, nitrification process, and the abundance of denitrification genes *nirS* and *nosZ*, but with no significant effect on AOB.^9,11,16,22^ Our results are consistent with previous findings except for AOA abundance, which may be due to our smaller sample size (n = 28, 4 studies) compared to others focused specifically on N-cycling genes.^22^ One benefit of biochar soil amendment is inhibiting N_2_O, a GHG 310 times more potent than CO_2_ and accumulating in the atmosphere at a rate of 2% per decade.^53^ Two thirds of global N_2_O emissions originate from soil, particularly agricultural soil with excessive N fertilizers. Based on 71 observations from 11 studies, we found that biochar could reduce cumulative N_2_O by 12.7%, although substantial within-study variation (80.8% of total heterogeneity) limited this effect’s significance. Earlier meta-analyses focused specifically on N_2_O emissions reported an overall 16%-50% N_2_O reduction by biochar.^5,19,20^ Further, our finding of reduced cumulative N_2_O despite the 40.8% increase in potential nitrification rate suggests strong inhibition of denitrification by biochar, which is supported by the observations of increased *nirS* (NO_2_^-^ → NO; 8.2%, *p* < 0.032) and *nosZ* (N_2_O → N_2_; 7.4%, *p* = 0.216). The main source of N_2_O is microbial NO reduction by diverse nitric oxide reductases (NOR).^51^ However, NOR genes are rarely measured. Future research could benefit from directly quantifying biochar’s effects on NOR genes and NOR activity.

Biochar enhanced the activity of two P-cycling enzymes despite lack of significance, and alkaline phosphatases showed greater enhancement than acid phosphatases (18.9% vs. 3.3%). Phosphatases hydrolyze phosphomonoesters and to a lesser extent phosphodiesters, releasing phosphate.^45^ Higher phosphatase activity could increase P availability, which is commonly seen in biochar treated soils.^54^ A prior meta-analysis reported similar results as ours but another one found insignificant decrease in acid phosphatase activity and significant increase in alkaline phosphatase activity.^9,11^ These discrepancies likely reflect variations in soil pH across studies, as the two phosphatases have different optimal pH ranges.^11,55,56^

### 3.3. Influence of Individual Moderators

The inclusion of a moderator reduced total non-sampling variance for all the responses (i.e., pseudo-*R*^2^ > 0), except for CFU (Table S6). But this also occasionally lowered the number of observations at a particular moderator level to <10 (in the SI). Overall, biochar characteristics, soil properties, and treatment protocols all influenced biochar’s effects on the soil microbiome (Figure 3; Figure S2).

**Figure 3.**
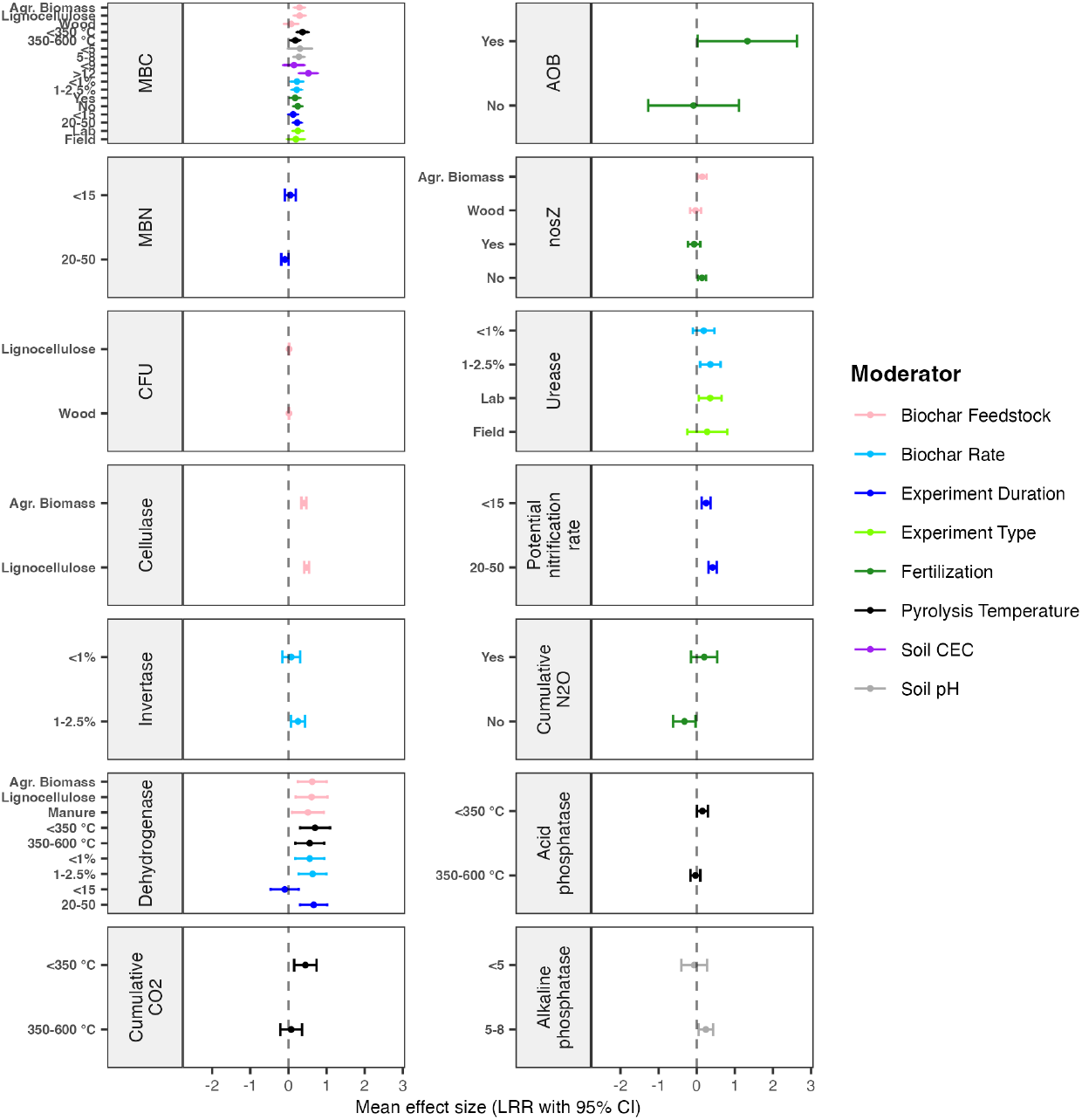
Mean effects of individual moderators on soil microbiome responses (weighted LRR ± 95% confidence interval). Data presented here had >10 observations at each moderator level. See the SI and Figure S3 for data with <10 observation at certain moderator levels. No significant effect was seen in β-glucosidase activity.

#### 3.3.1. Microbial Abundance and Diversity

Microbial biomass is often considered a soil health indicator and its increase is desired for soil funcitoning.^1^ Biochar’s positive effects on MBC were significantly influenced by all the moderators (*p* < 0.003 in omnibus test), except for soil C:N and soil zone (Figure 3). Biochar generated from agricultural biomass and lignocellulose had significant and similar effects (agricultural biomass: +34.39%, n = 67; lignocellulose: +33.81%, n = 33; both *p* < 0.001), while wood biochar had much smaller, non-significant effects (+7.38%, n = 99, *p* = 0.466). Biochar produced at low pyrolysis temperatures resulted in >2 times higher MBC increase than biochar produced at medium pyrolysis temperatures (≤350 °C: +45.11%, n = 24; 350-600 °C: +20.10%, n = 178; both *p* < 0.01), likely due to a higher amount of labile C and available nutrients in the former.^11^ MBC increases were slightly larger in acid soil (pH <5: +35.72%, n = 75, *p* = 0.061) than in neutral to alkaline soil (pH 5-8: +31.84%, n = 107, *p* < 0.001). High soil CEC (>12 cmol/kg) resulted in >4 times greater MBC increase than low soil CEC (<9 cmol/kg) (high CEC: +68.83%, n = 11, *p* < 0.001; low CEC: +15.56%, n = 46, *p* = 0.303). Low and medium biochar application rates yielded significant and similar MBC increases (<1% rate: +25.40%, n = 74; 1%-2.5% rate: +24.54%, n = 126; both *p* < 0.01). Higher MBC increases occurred without fertilization (no fertilizer: +27.83%, n = 162; fertilizer: +19.34%, n = 40; both *p* < 0.015). Longer experiment duration resulted in 2 times higher MBC increases (<15 days: +13.06%, n = 48, *p* = 0.070; 20-50 days: +26.09%, n = 150, *p* < 0.001). Slightly larger MBC increases occurred in the laboratory (+28.41%, n = 169, *p* < 0.001) than the field (+21.74%, n = 33, *p* = 0.100). These results were generally consistent with other reports of an 21.7% increase of MBC and greater MBC increases by biochar generated from agricultural biomass and lignocellulose than wood, or under lower than higher pyrolysis temperature.^11,23^ However, opposite to our findings, they observed smaller MBC increases in acidic soil than neutral soil, after medium than short experiment duration, and from laboratory than field experiments. These discrepancies are likely due to categorization criteria as acidic was defined as pH <6.5 and short duration was defined as up to 100 days in that study,^11^ as well as the distinguishment between measurement methods.^23^ Interestingly, we found that biochar increased rhizosphere MBC nearly 3 times more than bulk soil MBC (Figure S2). While the generality of this finding is limited by the small number of rhizosphere datasets (n = 2 for rhizosphere vs. n = 200 for bulk soil), it highlights potentially profound effects of biochar on the rhizosphere and warrants future research in this important yet overlooked direction.

For MBN, experiment duration was the only significant moderator (*p* = 0.039 in omnibus test) (Figure 3). Shorter duration increased MBN (+5.32%, n = 32) while longer duration decreased MBN (−8.76%, n = 80), but neither was significant. Our results, consistent with others,^11^ suggest limited effects of biochar on MBN. For bacterial and fungal PLFA, feedstock was the only significant moderator (*p* < 0.001) (Figure S2). Biochar generated from biosolids increased bacterial and fungal PLFA remarkably more than biochar made from lignocellulose or wood, despite of the small observation numbers (Figure S2).

Below-ground biodiversity is a key factor for maintaining the functioning of soil ecosystems and reduced microbial diversity could result in the decline of multiple soil functions including nutrient cycling and retention.^57^ However, soil health measurements often exclude diversity indicators, likely due to limited functional knowledge and lack of effective methods.^1^ Here observations at a particular moderator level were often less than 10 and occasionally, all from one study (Figure S2; Table S6). Nonetheless, we found substantially higher diversity increases in the following comparisons: biochar made from biosolids versus other feedstocks, rhizosphere versus bulk soil, with fertilizer versus without fertilizer, and in the field versus in the laboratory. We found no significant difference in diversity change among biochar application rates and marginally larger diversity increases from shorter versus longer experiments. These results are consistent with others. For example, one laboratory study found that biochar increased both prokaryotic and fungal diversity in the wheat rhizosphere, with greater diversity increases from medium (1-2%) than high (4%) application rate.^10^ One meta-analysis focused specifically on microbial diversity reported greater Shannon index increases by biochar made from manure and sludge versus other feedstocks.^23^ However, the same study found larger bacterial diversity increases from field versus laboratory experiments, but larger fungal diversity increases from laboratory versus field experiments.

#### 3.3.2. C, N, P-Cycling Functions and Processes

For C-cycling enzymes and cumulative CO_2_, biochar feedstock, pyrolysis temperature, soil CEC, fertilizer addition, biochar application rate, and experiment duration were significant moderators (*p* < 0.035 in omnibus test) (Table S6). Significant moderators for cellulase activity included biochar feedstock and experiment duration. Lignocellulose biochar promoted cellulase activity more than biochar generated from agricultural biomass (agricultural biomass: +50.11%, n = 20; lignocellulose: +61.22%, n = 20; both *p* < 0.001); longer experiments resulted in larger increases than shorter experiments (<15 days: +42.14%, n = 8; 20-50 days: +59.14%, n = 32; both *p* < 0.001) (Figure 3; Figure S2). Significant moderators for invertase activity included biochar feedstock, pyrolysis temperature, fertilizer addition, and biochar application rate. In general, larger invertase activity increases occurred by biochar made from agricultural biomass and at medium pyrolysis temperature (agricultural biomass: +31.83%, n = 19; 350-600 °C: +28.65%, n = 25; both *p* < 0.002), with fertilizers, and at medium biochar application rate (fertilizer: +49.99%, n = 4; 1%-2.5% rate: +28.64%, n = 17; both *p* < 0.01) (Figure 3; Figure S2). Significant moderators for dehydrogenase activity were biochar feedstock, pyrolysis temperature, soil CEC, application rate, and experiment duration. Greater dehydrogenase activity increases occurred with agricultural biomass or lignocellulose feedstock (agricultural biomass: +86.90%, n = 69; lignocellulose: +84.00%, n = 23; manure: +67.03%, n = 36; all *p* < 0.02), low pyrolysis temperature (≤350 °C: +101.14%, n = 42; 350-600 °C: +74.63%, n = 86; both *p* < 0.005), high soil CEC (>12 cmol/kg: +78.00%, n = 6; <9 cmol/kg: +28.96%, n = 40; both *p* < 0.001), medium biochar application rate (<1% rate: +88.71%, n = 52; 1%-2.5% rate: +74.69%, n = 76; both *p* < 0.005), and longer experiment duration (<15 days: -9.17%, n = 32, *p* < 0.001; 20-50 days: +93.77%, n = 96, *p* = 0.609) (Figure 3; Figure S2). Another meta-analysis also found low pyrolysis temperature resulting in significantly higher dehydrogenase activity, although it did not consider fixed effect of moderators and had small observation sizes for many moderators.^11^ For cumulative CO_2_, pyrolysis temperature was the only significant. Low pyrolysis temperature resulted in greater increases in cumulative CO_2_ (≤350 °C: +56.03%, n = 12, *p* < 0.004; 350-600 °C: +7.64%, n = 26, *p* = 0.602) (Figure 3). This trend is consistent with low-temperature biochar’s stronger effect on soil microbial metabolic state, exemplified by dehydrogenase activity.

For N-cycling genes (*amoA* of AOB, *nirS, nosZ*), enzyme (urease), and processes (potential nitrification rate, cumulative N_2_O), biochar feedstock, pyrolysis temperature, soil C:N ratio, fertilizer addition, application rate, experiment duration, and field or laboratory were significant moderators (*p* < 0.05 in omnibus test) (Table S6). No significant moderator was identified for AOA *amoA* or *narG* (NO_3_^-^ → NO_2_^-^) (Table S6). For AOB, fertilization was the only significant moderator. Together with fertilization, biochar increased AOB by nearly 3 times (no fertilizer: -7.75%, n = 48, *p* = 0.892; fertilizer: +277.45%, n = 13, *p* < 0.05) (Figure 3). Significant moderators for *nirS* included feedstock and experiment duration. Wood biochar and shorter duration resulted in larger *nirS* increases (wood: +13.54%, n = 20, *p* < 0.004; <15 days: +33.07%, n = 4, *p* <0.001) (Figure S2). Significant moderators for *nosZ* were Biochar feedstock and fertilization. Biochar generated from agricultural biomass increased *nosZ* while wood biochar reduced *nosZ* abundance (agricultural biomass: +15.63%, n = 16, *p* = 0.016; wood: - 2.97%, n = 20, *p* = 0.673) (Figure 3). Fertilization revoked biochar’s positive effect on *nosZ* (no fertilizer: +15.00%, n = 23, *p* = 0.013; fertilizer: -6.46%, n = 13, *p* = 0.407). Significant moderators of urease activity included biochar feedstock, pyrolysis temperature, soil zone, biochar application rate, fertilization, and field or laboratory experiments; however, only feedstock reduced total non-sampling variance and the observation number for certain moderators was less than 10 (Figure S2). Urease activity increases were higher under medium application rate (<1% rate: +20.09%, n = 14, *p* = 0.196; 1%-2.5% rate: +43.04%, n = 58, *p* < 0.01) and in the laboratory (laboratory: +42.32%, n = 60, *p* = 0.025; field: +31.46%, n = 14, *p* = 0.301) (Figure 3), by biochar made from lignocellulose and at low pyrolysis temperature, in the rhizosphere, and without fertilization (Figure S2). For potential nitrification rate, experiment duration was the only significant moderator. Longer experiments increased potential nitrification rate more than shorter experiments (<15 days: +28.26%, n = 15; 15-20 days: +52.09%, n = 18; both *p* < 0.001) (Figure 3). For cumulative N_2_O emissions, significant moderators included feedstock, soil C:N, and fertilization, but certain moderator levels had <10 observations (in the SI). In general, fertilization curtailed biochar’s inhibition of cumulative N_2_O emissions (no fertilizer: -27.43%, n = 41, *p* = 0.293; fertilizer: 21.65%, n = 30, *p* = 0.343) (Figure 3). All feedstocks but wood reduced cumulative N_2_O emissions and inhibition was significant in soils with lower C:N (Figure S2).

For P-cycling enzymes, biochar feedstock, pyrolysis temperature, and soil pH were significant moderators (*p* < 0.05 in omnibus test) (Table S6). Acid phosphatase activity was significantly increased by biochar made at low pyrolysis temperature (≤350 °C: +16.16%, n = 16, *p* < 0.045; 350-600 °C: -3.24%, n = 70, *p* = 0.610) (Figure 3). Alkaline phosphatase activity was enhanced by biochar made from agricultural biomass more than other feedstocks (agricultural biomass: +84.92%, n = 4, *p* < 0.001), and only in neutral to alkaline soils (pH <5: -6.36%, n = 24, *p* = 0.694; pH 5-8: +27.29%, n = 14, *p* < 0.017) (Figure S2).

### 3.4. Importance of Moderators in Precision Biochar Applications

Feedstock, pyrolysis temperature, soil pH, fertilization, biochar application rate, and experiment duration were the most important predictors for microbial responses (Figure 4; Table S7) and most frequently included in the final models (Table S8). Feedstock was an important predictor for bacterial PLFA, β-glucosidase, and cumulative N_2_O emissions. Pyrolysis temperature was an important predictor for dehydrogenase, cumulative CO_2_ emissions, and acid phosphatase activity. Soil pH was an important predictor for fungal PLFA, Chao1 diversity, and acid phosphatase activity. Biochar application rate was an important predictor for MBC, urease activity, and potential nitrification rate. Fertilization was an important predictor for AOB abundance and cumulative N_2_O emissions. Experiment duration was an important predictor for potential nitrification rate and acid phosphatase activity.

**Figure 4.**
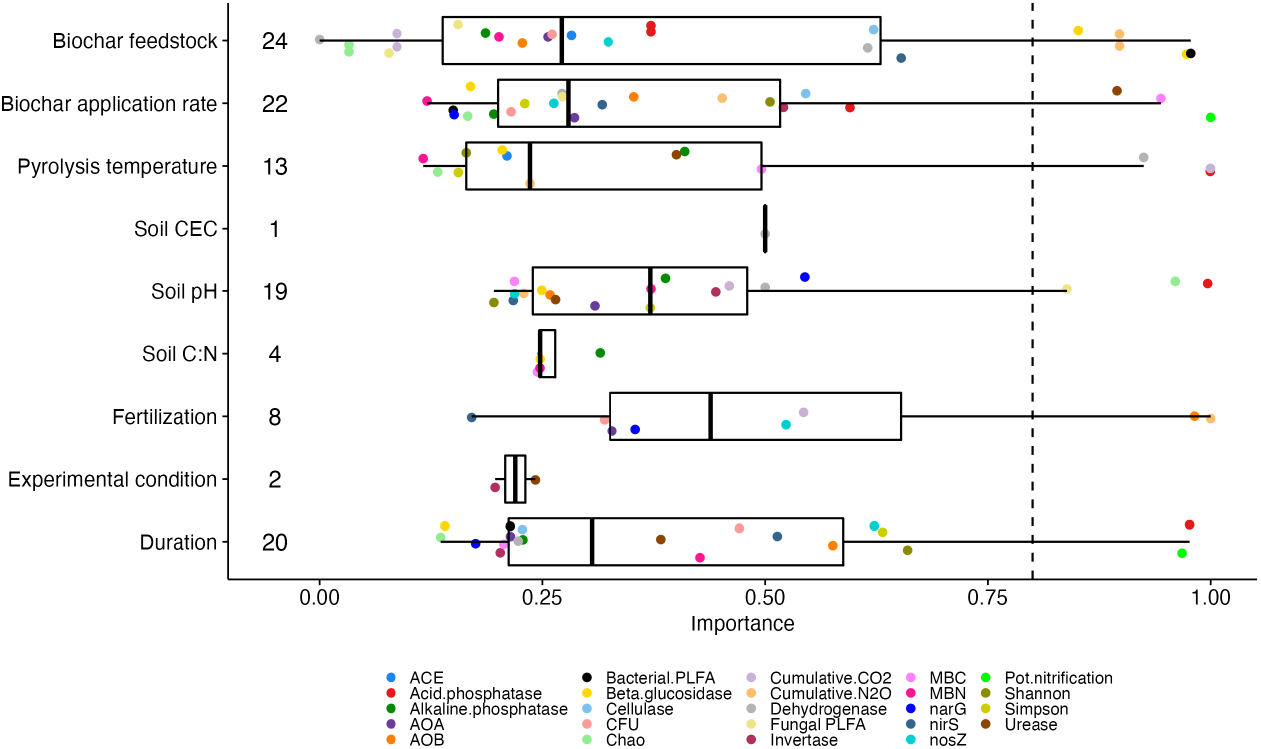
AICc-based predictor importance for the 24 soil microbiome responses included in this study. The number behind a predictor is the number of models containing that predictor.

To achieve a specific soil amendment goal, precision biochar application could be implemented with careful control of these factors and multi-criteria decision analysis could be conducted to prioritize soil management outcomes. First, biochar production parameters can be tuned to elicit the desired microbial responses. Feedstock and pyrolysis temperature are major controls of biochar physicochemical properties, such as porosity, specific surface area, crystallinity, fixed carbon, volatile and ash content, elemental composition, surface functional group, and recalcitrance.^58–61^ Reduction of N_2_O emissions can be better achieved with biochar made from lignocellulose feedstock. Biochar made from agricultural biomass is also promising, as it could simultaneously boost *nirS* and *nosZ* genes that mediate the conversion of NO_2_^-^ to NO and N_2_O to N_2_, respectively. Biochar made at lower pyrolysis temperatures is preferred for acid phosphatase and dehydrogenase that can increase soil CEC and thus nutrient retention; however, CO_2_ reduction requires biochar made at higher temperature. Therefore, an optimal amendment plan could switch biochar generated at multiple temperatures during the growth season to reach both goals. Second, biochar application rate and duration could be optimized to invoke the best outcomes. Relatively higher application rate could enhance urease and nitrification rate more, but may reduce acid phosphatase activity. Optimal application rates can be determined based on the needs of different crops. Moreover, our model showed that N_2_O inhibition was stronger in shorter durations, suggesting that even with the optimal biochar, repeated biochar amendment may be needed for best mitigation outcome.

### 3.5. Knowledge Gaps and Future Directions

We identified several critical knowledge gaps hindering the realization of biochar-based sustainable soil management. First, mechanistic understanding of biochar-induced soil microbiome alterations is still largely unknown. Multi-omics holds the promise to shed light on genome-to-phenome changes in the soil microbiome.^10,62^ The rhizosphere microbiome at the interface between plant roots and soil deserves particular attention as it links below-ground processes to above-ground plant development and health. However, we only found three studies focused on rhizosphere microbiome responses. Multi-omics may also offer an in-depth understanding of N_2_O inhibition by biochar, where multiple mechanisms have been proposed, including reduced denitrification due to aeration, adsorption of reactive N intermediates, shifted denitrification stoichiometry due to pH increase, and more complete denitrification by denitrifiers due to available labile C or biochar-facilitated electron transfer.^20,63–65^ Biochar can influence multiple pathways in microbially mediated N cycling, which can eventually shift the N_2_O/(N_2_+N_2_O) ratio.^22,63^ The expression and activity of several reductases (NIR, NOR, NOS) requires particular attention. Second, biochar varies significantly in physicochemical properties and without characteristic data, experimental results cannot be appropriately compared. Specifically, biochar’s redox activity is important to many microbially mediated transformations in the soil, but it is not widely reported. Lignocellulose is an essential source of redox moieties such as electron-donating phenolic moieties and electron-accepting quinones. Pyrolysis temperature influences the surface density of these moieties and the ordering of carbon structures, thus affecting charging and discharging of surface functional groups and direct electron transfer, two electron flow pathways seen in biochar.^66,67^ Future efforts should address, with tools such as machine learning, how to leverage biochar redox characteristics to evoke desired microbial responses such as N_2_O reduction to N_2_. Ultimately, this work advances our understanding of the soil microbiome’s responses to biochar amendment and highlights purpose-driven biochar fabrication and application as a promising strategy for sustainable agriculture.

## Supporting Information

Detailed materials and methods (e.g., all primary studies used in this meta-analysis, best model selection procedure), additional results and discussion (e.g., reduction of total non-sampling variance from random-effects models), figures (an increasing number of biochar studies over the past decades, significant single categorical moderators, funnel plots for global effect sizes of individual soil microbial responses before model selection and in their final models), tables (keyword combinations, a complete list of response variables extracted from all primary studies, complete list of predictor variables used in this study, full results of three-level random-effects models) (PDF).

Raw data extracted from the 61 primary studies, full summary of statistics of the three-level mixed-effects models including one single categorical moderator, importance of individual predictors to soil microbial responses, full summary of statistics of the final three-level mixed-effects models, assessment of publication bias in the three-level random-effects models including one single categorical moderator, assessment of publication bias in the final best three-level random-effects models including multiple moderators (XLSX).

## Acknowledgement

This work was supported by Idaho State University through a Developing Collaborative Partnerships Grant. Y. You also received support from the Center for Advanced Energy Studies (CAES), a research, education, and innovation consortium consisting of Idaho National Laboratory and the public research universities of Idaho. Patricia Kerner is grateful to Idaho State University Center for Ecological Research and Education. We are grateful to all the researchers whose data contributed to this meta-analysis.

