## Supplemental Information for "Microbial Responses to Biochar Soil Amendment and Influential Factors: A Three-level Meta-analysis"

Number of pages: 32
Number of tables: 10
Number of figures: 4

### S1. Supplementary Materials and Methods

#### S1.1. Systematic Literature Review

During the past decades, biochar soil amendment has received increasing attention from the research community due to its beneficial effects on soil health, crop production, and environmental sustainability (Figure S1).[1] Here we conducted a systematic review using the keyword "biochar" paired with soil microbial response terms in the Web of Science (keywords and keyword combinations in Table S1). In brief, we searched for both bacterial and fungal responses, ranging from abundance and diversity to community structure and composition and to enzyme activity and community function. After literature screening as detailed in the PRISMA diagram (Figure 1), a total of 61 studies were retained.[2–62] The 61 studies included experiments conducted in the field (14 studies) or laboratory, spanning a wide range of climatic regions. A total of 3899 pairwise observations were obtained from these studies and included in the meta-analysis.

#### S1.2. Three-level Meta-analytical Model

The estimate of study variance ($\hat{\sigma}^2$) was calculated using the reported variance of the mean of a response variable (Eq. 2 in the main text). When only standard errors were reported, sample standard deviations were back calculated. If a study did not report variance, the information was obtained from the authors.

For each response variable, a three-level random-effects model was used, assuming random effects at different levels and independent sampling error (Eq. 3 in the main text):[63]

$$y_{ij} = \beta_0 + u_{(2)ij} + u_{(3)j} + e_{ij} \quad \text{(S1)}$$

Where $y_{ij}$ is the *i*th effect size in the *j*th study; $\beta_0$is the average population effect; $Var(e_{ij}) = v_{ij}$ ($e_{ij}$ being the sampling error in the *i*th effect size in the *j*th study); $Var(u_{(2)ij}) = \tau^2_{(2)}$ is the within-study heterogeneity between effect sizes, $Var(u_{(3)j}) = \tau^2_{(3)}$ is the between-study heterogeneity. We assumed that errors and random effects were normally distributed with a mean of 0 and independent:

$$Cov(u_{(2)ij}, u_{(3)j}) = Cov(u_{(2)ij}, e_{ij}) = Cov(u_{(3)j}, e_{ij}) = 0 \quad \text{(S2)}$$

We used Cochran's Q to test the significance of heterogeneity.[64] We used the statistic $I^2$ developed by Higgins and Thompson (2002) to estimate the proportion of variation in effect sizes explained by level-2 or level-3 variance, with total heterogeneity being the sum of both. The statistic $I^2$ is a transformation of $H$:[65]

$$I^2 = \frac{H^2 - 1}{H^2} \quad \text{(S3)}$$

where $H$ is the square root of the $\chi^2$ heterogeneity statistic divided by its degrees of freedom. Thus $I^2$ describes the proportion of total variation in effect estimates across studies due to heterogeneity.

Multicollinearity of candidate predictors was evaluated by including all predictors in a model and using variance inflation factors (VIF). If there was high correlation (VIF > 5), the predictor with the highest VIF was removed from the model, and the model was checked once again. This process was repeated until there was no correlation among predictors. The R package “glmulti” was used for automated model selection from all possible combinations of uncorrelated predictors. For each response variable, the importance of a moderator was computed as the sum of model probabilities (i.e., weights) for the models that involved the moderator (i.e.., model-averaged moderator importance). A moderator was considered as important if its model-averaged importance was >0.8. The model with lowest corrected Akaike information criterion (AICc) value was considered as the best model.

#### S1.3. Publication Bias

Missing data is a critical issue in meta-analysis. Techniques for assessing missing data in conventional meta-analysis include the trim-and-fill method (i.e., funnel plot), Egger’s linear regression test, Kendall’s tau rank correlation test, and Rosenthal’s fail-safe test (i.e., fail-safe number (FSN) analysis). However, these approaches have not been comprehensively evaluated in multi-level meta-analysis, leaving it difficult to select the most appropriate method for handling missing data in this study.[66] Consequently, we applied all the three aforementioned techniques in this study. Potential influence of publication bias on the validity of global effect sizes was first analyzed using funnel plots, and the significance of funnel plot asymmetry was tested using Egger’s linear regression test.[67] When significant asymmetry was observed, the fail-safe number (FSN) analysis was conducted using the Rosenberg method.[68] Notably, FSN can vary substantially and there is no standard cutoff for what is considered a large FSN.[69]

### S2. Supplementary Results and Discussion

#### S2.1. Global Effect Sizes

Biochar had positive effects on microbial abundance indices CFU, MBC, bacterial PLFA, and fungal PLFA, but negative effects on MBN. Such effects were significant for CFU and MBC. Biochar effects on microbial diversity indices were all positive and ranged from +3.0% to +8.4%, even though they were insignificant.

Biochar had positive effects on all the C-cycling related responses. Such effects were significant for the activity of cellulases that decompose cellulose and related polysaccharides, dehydrogenases that are involved in oxidation-reduction of organic compounds, and invertases that hydrolyze sucrose, but insignificant for the activity of β-glucosidases that catalyze the final step of cellulose hydrolysis by converting cellobiose to glucose.

Biochar had positive effects on the N-cycling related response AOB, *narG*, *nirS*, *nosZ*, potential nitrification rate, and urease activity. Such effects were significant for the abundance of nitrite reductase gene *nirS*, the activity of ureases that hydrolyze urea, and potential nitrification rate. Biochar had negative effects on two N-cycling related response AOA and cumulative $N_2O$, although such effects were insignificant.

Biochar had positive effects on the activity of acid phosphatases and alkaline phosphatases, with alkaline phosphatase activity showing stronger response, although neither was significant.

**S2.2. Within- and Between-study Variance in the Three-level Random-effects Model**

Cochran's Q test suggests that between-study variation was significant ($p < 0.05$) for all the response variables except for ACE (Table S5). Considering the limitation of Cochran's Q test when handling a small number of studies, we also calculated the statistic $I^2$ that is independent of the number of studies (Table S5). For response variables other than CFU and ACE, models explained 77.7%-99.9% of total heterogeneity (level-2 $I^2$ + level-3 $I^2$).

Within-study variance (level-2 $I^2$) accounted for >80% of total heterogeneity for CFU, ACE, cellulase activity, AOA abundance, *nirS* and *nosZ* gene abundance, cumulative $N_2O$, and potential nitrification rate. In particular, CFU, cellulase activity, and potential nitrification rate had a large proportion of within-study variance partially because all the observations were from a single study (i.e., zero level-3 $I^2$). On the other hand, between-study variance (level-3 $I^2$) accounted for >86% of total heterogeneity for bacterial PLFA, the diversity index Simpson, Shannon and Chao1, urease activity, and cumulative $CO_2$.

**S2.3. Influence of Individual Moderators**

For some microbial responses, the inclusion of a moderator decreased the number of observations to <10 at a particular moderator level. Consequently, the robustness of the findings for these microbial responses and moderators may be limited (Table S6).

**S2.3.1. Microbial Abundance and Diversity**

Biochar generated from lignocellulose and wood had similarly marginal and significant effects on bacterial CFU (lignocellulose: +1.95%, n = 16; wood: +1.55%, n = 16; $p < 0.001$) (Figure 3). This was also the case for biochar application rate (1% rate: +1.72%, n = 24; 1-2.5% rate: +1.83%, n = 8; both $p < 0.005$), fertilizer addition (no fertilizer: +1.57%, n = 24; fertilizer: +2.31%, n = 8; both $p < 0.001$), and experiment duration (<15 days: +1.69%, n = 8; 15-20 days: +1.77%, n = 24; both $p < 0.007$), although the observation number at certain moderator levels dropped to less than 10 (Figure S2).

Biochar's positive effects on MBC were significantly influenced by all the moderators ($p <$ 0.003 in omnibus test), except for soil C:N and soil zone (Figure 3). Soil CEC and pyrolysis temperature explained the highest within- and between-study variance, respectively (40% and 74% of total $I^2$) (Table S6). But only biochar feedstock and soil CEC reduced total non-sampling variance of the random-effects model (pseudo-$R^2$ = 13% and 40%, respectively). Interestingly, biochar increased rhizosphere MBC nearly 3 times more than bulk soil MBC (rhizosphere: +70.37%, n = 2, $p = 0.071$; bulk: +24.83%, n = 200, $p < 0.001$) (Figure S2). While the generality of this finding is limited by the small number of rhizosphere data, it highlights potentially profound effect of biochar on the rhizosphere and warrants future research in this direction.

For MBN, experiment duration was the only significant moderator ($p = 0.039$ in omnibus test) and within-study variance explained 83% of total $I^2$. The inclusion of this moderator reduced total non-sampling variance of the random-effects model by 40% (Table S6). Shorter duration increased MBN (+5.32%, n = 32) while longer duration decreased MBN (-8.76%, n = 80), but neither was significant (Figure 3).

For bacterial and fungal PLFA, feedstock was the only significant moderator ($p < 0.001$ in omnibus test), reducing total non-sampling variance by 87% and 56%, respectively (Table S6). Within-study variance accounted for 97.9-99.2% of total $I^2$. Biochar generated from biosolids resulted in >170 times greater bacterial PLFA increase than biochar generated from wood (biosolids: +149.90%, n = 4, $p < 0.001$; wood: +0.86%, n = 23, $p = 0.830$) (Figure S2). Biochar generated from biosolids resulted in almost 4 times higher fungal PLFA than biochar generated from lignocellulose (biosolids: +81.43%, n = 4, $p < 0.001$; lignocellulose: +21.98%, n = 3, $p = 0.208$), whereas biochar generated from wood had no significant effect on fungal PLFA (-0.69%, n = 23, $p = 0.899$) (Figure S2).

For diversity indices ACE, Chao1, Shannon and Simpson, biochar feedstock, soil zone, biochar application rate, fertilizer addition, experiment duration, and field or laboratory were significant moderators ($p < 0.025$ in omnibus test), although observations at a particular moderator level were often less than 10 and occasionally, all came from a single study. For ACE, fertilizer addition was the only significant moderator ($p < 0.025$ in omnibus test) and reduced total non-sampling variance by 30%. Slightly higher ACE increases occurred with fertilizer addition (no fertilizer: +1.33%, n = 19, $p = 0.489$; fertilizer: +6.29%, n = 4, $p < 0.009$) (Figure S2).

For Chao1, soil zone and experiment type were two significant moderators ($p < 0.001$ in omnibus test), each reducing total non-sampling variance by 79%. Increases in Chao1 were >50 times higher in the field (+47.81%, n = 1, $p < 0.001$) than in the laboratory (+0.93%, n = 46, $p = 0.628$) (Figure S2). Rhizosphere soil showed >50 times higher Chao1 increases than bulk soil (rhizosphere: +43.81%, n = 2, $p < 0.001$; bulk: +0.80%, n = 45, $p = 0.678$). However, field observations and rhizosphere data were extremely limited; thus the generality of this finding needs further validation.

For Shannon index, biochar feedstock, experiment duration, and experiment type were three significant moderators ($p < 0.015$ in omnibus test). The inclusion of feedstock or experiment type reduced the total non-sampling variance by >98%, whereas experiment duration did not. Biochar made from biosolids resulted in remarkably higher Shannon increases than biochar made from wood, agricultural biomass, or lignocellulose (biosolids: +69.07%, n = 4, $p < 0.001$; wood: +2.71%, n = 16, $p = 0.047$; lignocellulose: -0.04%, n = 10, $p = 0.980$; agricultural biomass: -0.48%, n = 30, $p = 0.636$) (Figure S2). Shorter and longer experiment duration had similar and insignificant effects on Shannon index (<15 days: +8.52%, n = 6, $p = 0.133$; 15-20 days: +6.01%, n = 50, $p = 0.276$). More than 100 times higher Shannon increases occurred in the field (+69.07%, n = 4, $p < 0.001$) than in the laboratory (+0.53%, n = 56, $p = 0.421$). It should be noted that all the field data were from one primary study which used biosolids biochar. This limits the generality of these findings and warrant more field research using biochar made from other feedstocks.

For Simpson index, feedstock, biochar application rate, and experiment type were the three significant moderators ($p < 0.001$ in omnibus test). The inclusion of biochar feedstock or experiment type reduced the total non-sampling variance by 99.7% and 98.7%, respectively, whereas biochar application rate did not. Biochar generated from biosolids resulted in remarkably higher Simpson increases than biochar generated from wood, lignocellulose, or agricultural biomass (biosolids: +103.50%, n = 4, $p < 0.001$; wood: +2.79%, n = 16, $p = 0.011$; lignocellulose: -0.13%, n = 2, $p = 0.994$; agricultural biomass: -0.89%, n = 16, $p = 0.228$) (Figure S2). Low and medium biochar application rate yielded similar and insignificant Simpson increases (1% rate: +8.11%, n = 7, $p = 0.300$; 1-2.5% rate: 8.43%, n = 31, $p = 0.282$). More than

100 times higher Simpson increases occurred in the field (+103.50%, n = 4, $p < 0.001$) than in the laboratory (+0.56%, n = 34, $p = 0.598$). However, all the field data were from one primary study which used biosolids biochar. More field research using biochar made from other feedstocks is required to validate the findings here.

#### S2.3.2. C, N, P-Cycling Functions and Processes

##### S2.3.2.1. C Cycling

For C-cycling enzymes (cellulase, dehydrogenase, and invertase), biochar feedstock, pyrolysis temperature, soil CEC, biochar application rate, fertilization, and experiment duration were significant moderators ($p < 0.05$ in omnibus test) (Figure 3 and Figure S2). For cellulase activity, feedstock and experiment duration were significant moderators but they only reduced total non-sampling variance of the random-effects model marginally (pseudo-$R^2$ < 9%) (Table S6). Within-study variance accounted for >91% of total $I^2$.

For invertase activity, feedstock and pyrolysis temperature contributed more to within-study heterogeneity (64% and 59% of total $I^2$, respectively), while fertilization and biochar application rate contributed more to between-study heterogeneity (54% and 63% of total $I^2$, respectively) (Table S6). The inclusion of biochar feedstock or pyrolysis temperature reduced total non-sampling variance of the random-effects model by 18% and 16%, respectively, while the inclusion of fertilization or biochar application rate did not reduce total non-sampling variance. Biochar generated from agricultural biomass and wood feedstocks had similar effects on invertase activity and resulted in larger increases than lignocellulosic feedstocks (agricultural biomass: +31.83%, n = 19, $p < 0.002$; wood: +21.66%, n = 6, $p = 0.063$; lignocellulose: +4.15%, n = 6, $p = 0.701$) (Figure S2). Medium pyrolysis temperatures resulted in higher increases in invertase activity than low pyrolysis temperature (<350 °C: +4.35%, n = 6, $p = 0.700$; 350-600 °C: +28.65%, n = 25, $p < 0.002$). Fertilizer addition resulted in greater increases compared to the absence of fertilizer (fertilizer: +49.99%, n = 4, $p = 0.008$; no fertilizer: +18.71%, n = 27, $p = 0.027$).

For dehydrogenase activity, significant moderators for dehydrogenase activity were biochar feedstock, pyrolysis temperature, soil CEC, application rate, and experiment duration. Feedstock, pyrolysis temperature and biochar application rate explained both within- and between-study heterogeneity, each attributing to ~50% of total $I^2$ and together accounted for >99.9% of total $I^2$ (Table S6). Soil CEC was more responsible for within-study variance (74% of total $I^2$) while experiment duration more explained between-study variance (71% of total $I^2$).

The inclusion of soil CEC or experiment duration as a single moderator reduced total non-sampling variance by 98% and 29%, respectively.

For cumulative $CO_2$, pyrolysis temperature was the only significant moderator, reducing 47% of total non-sampling variance (Table S6). Low pyrolysis temperature resulted in greater increases in cumulative $CO_2$ than medium pyrolysis temperature (≤350 °C: +56.03%, n = 12, $p <$ 0.004; 350-600 °C: +7.64%, n = 26, $p = 0.602$).

**S2.3.2.2. N Cycling**

For N-cycling related genes (*amoA* of AOB, *nirS*, *nosZ*), enzyme (urease) and processes (potential nitrification rate, cumulative $N_2O$), biochar feedstock, pyrolysis temperature, soil C:N ratio, fertilizer addition, application rate, experiment duration, and field or laboratory were significant moderators ($p < 0.05$ in omnibus test) (Figure 3 and Figure S2). No significant moderator was identified for *amoA* of AOA or *narG* ($NO_3^- \rightarrow NO_2^-$) (Table S6). Fertilization was the only significant moderator for AOB *amoA* and reduced 6% of total non-sampling variance (Table S6).

Biochar feedstock and experiment duration were significant moderators for *nirS* gene abundance and within-study variance explained >99% of total $I^2$(Table S6). The inclusion of these moderators reduced total non-sampling variance of the random-effects model by 12% and 66%, respectively. Biochar generated from wood increased *nirS* more than agricultural biomass (agricultural biomass: +2.69%, n = 8, $p = 0.541$; wood: +13.54%, n = 20, $p < 0.004$). Short experiments resulted in greater *nirS* increases than longer experiments (<15 days: +33.07%, n = 4, $p <0.001$; 20-50 days: +2.89%, n = 24, $p = 0.198$) (Figure S2).

Biochar feedstock and fertilization were significant moderators for *nosZ* abundance and the inclusion of them reduced 12% and 13% of total non-sampling variance, respectively (Table S6). Within-study variance accounted for >99% of total heterogeneity.

For urease activity, significant moderators included biochar feedstock, pyrolysis temperature, soil zone, biochar application rate, fertilization, and field or laboratory. However, only feedstock reduced total non-sampling variance of the random-effects model by 33% and the number of observations for certain moderators was less than ten (Table S6). Larger urease activity increases occurred with lignocellulose biochar (agricultural biomass: +22.83%, n = 60, $p = 0.086$; lignocellulose: +118.87%, n = 8, $p < 0.001$; wood: +2.81%, n = 6; all $p = 0.840$), low pyrolysis temperature (≤350 °C: +92.29%, n = 7, $p < 0.015$; 350-600 °C: +26.42%, n = 66; both $p = 0.130$), the rhizosphere (rhizosphere: +62.71%, n = 2, $p = 0.282$ ; bulk soil: +37.57%, n = 72,

$p < 0.025$), and no fertilizer (no fertilizer: +40.54%, n = 69, $p < 0.01$; fertilizer: +24.85%, n = 5, $p = 0.148$) (Figure S2).

Experiment duration was a significant moderator for potential nitrification rate (Figure 3). The inclusion of this moderator reduced 11% of total non-sampling variance of the random-effects model (Table S6).

For cumulative $N_2O$ emissions, significant moderators included feedstock, soil C:N, and fertilization. The inclusion of these moderators reduced total non-sampling variance by 14%, 67%, and 12%, respectively. Except for wood, all feedstocks inhibited cumulative $N_2O$ emissions (agricultural biomass: -36.97%, n = 5, $p = 0.164$; lignocellulose: -31.05%, n = 26, $p < 0.015$; wood: +24.01%, n = 36, $p = 0.089$; biosolids: -61.13%, n = 2, $p = 0.075$; manure: -16.08%, n = 2, $p = 0.738$) (Figure S2). Inhibition of cumulative $N_2O$ emissions occurred in soils with lower or higher C:N, but this effect was significant only under lower soil C:N (C:N <12: -32.30%, n = 8, $p = 0.047$; C:N >13: -32.35%, n = 18, $p = 0.073$).

**S2.3.2.3. P Cycling**

For P-cycling related enzymes (acid and alkaline phosphatase), biochar feedstock, pyrolysis temperature, and soil pH were significant moderators ($p < 0.05$ in omnibus test) (Figure 3 and Figure S2). Pyrolysis temperature was a significant moderator for acid phosphatase activity but it did not reduce total non-sampling variance of the random-effects model (Table S6).

For alkaline phosphatase activity, feedstock and soil pH were significant moderators. The inclusion of feedstock or soil pH reduced 37.5% and16% of total non-sampling variance in the random-effects model, respectively (Table S6). Biochar generated from agricultural biomass increased alkaline phosphatase activity the most, while lignocellulose and wood biochar had similar effects (agricultural biomass: +84.92%, n = 4, $p < 0.001$; lignocellulose: +7.81%, n = 18, $p = 0.324$; wood: +5.77%, n = 16, $p = 0.512$) (Figure S2).

**S2.4. Importance of Moderators in Precision Biochar Applications**

In a meta-analysis, moderators that account for a larger amount of variance between studies are more important because they help clarify which factors are driving the global effect sizes observed. Feedstocks used to generate biochar range from woody materials to agricultural biomass to animal manure or biosolids from wastewater treatment. These starting materials, along with pyrolysis temperature, mostly govern the resulting biochar's physicochemical characteristics.

#### S2.4.1. Microbial Abundance and Diversity

Biochar application rate was an important predictor (importance > 8) for MBC and the best mixed-effects model of MBC also included pyrolysis temperature (n = 43, $p = 0.003$ in omnibus test) (Tables S7 and S8). However, the best model did not reduce the total non-sampling variance of the random-effects model (pseudo-$R^2$ = 0). For bacterial PLFA, biochar feedstock was the only important predictor included in the final model and largely reduced the total non-sampling variance of the random-effects model (n = 18, $p < 0.001$ in omnibus test, pseudo-$R^2$ = 83%). The final model for fungal PLFA only included soil pH as the only important moderator which largely reduced the total non-sampling variance of the random-effects model (n = 15, $p < 0.001$ in omnibus test, pseudo-$R^2$ = 62%). The final MBN model and CFU model did not include any moderator.

The final models of all the microbial diversity indices, except for ACE, contained moderators (Tables S7 and S8). The final Chao1 model included soil pH as an important predictor (n = 20, $p < 0.001$ in omnibus test). However, the inclusion of soil pH did not reduce the total non-sampling variance of the random-effects model (pseudo-$R^2$ = 0). Biochar application rate and experiment duration were included in the best model of Shannon diversity but neither was important (n = 25, $p = 0.013$ in omnibus test, pseudo-$R^2$ =0.5%). The best Simpson model included experiment duration as the only but non-important predictor (n = 23, $p$ = 0.038 in omnibus test, pseudo-$R^2$ = 0%).

#### S2.4.2. C, N, P-Cycling Functions and Processes

##### S2.4.2.1. C Cycling

The best models of the C-cycling related responses all contained moderators (Tables S7 and S8). The best model of β-glucosidase had feedstock (lignocellulose or wood) as an important predictor (n = 24, $p < 0.001$ in omnibus test, pseudo-$R^2$ = 88.9%). The best model of cellulase included feedstock and biochar application rate as two non-important predictors (n = 40, $p = 0.036$ in omnibus test, pseudo-$R^2$ = 19.3%). The best model of dehydrogenase included feedstock, pyrolysis temperature, and soil CEC as moderators (n = 46, $p < 0.001$ in omnibus test, pseudo-$R^2$ = 98.9%), but only pyrolysis temperature was important. The best model of invertase included biochar application rate as the only but non-important predictor (n = 24, $p$ = 0.037 in omnibus test, pseudo-$R^2$ = 53.4%). Pyrolysis temperature and fertilization were included in the final model of cumulative $CO_2$ emissions (n = 34, $p < 0.001$ in omnibus test, pseudo-$R^2$ = 49.1%), but only pyrolysis temperature was an important predictor.

**S2.4.2.2. N Cycling**

Except for AOA, the final models of the N-cycling related responses all contained moderators (Tables S7 and S8). The best AOB model contained fertilization as one important predictor and experiment duration as a non-important predictor (n = 61, $p < 0.001$ in omnibus test, pseudo-$R^2$ = 24.8%). The best model of *narG* included soil pH as a single, non-important predictor (n = 18, $p = 0.011$ in omnibus test, pseudo-$R^2$ = 54.0%). The final model of *nirS* included feedstock and experiment duration as two non-important predictors (n = 28, $p = 0.015$ in omnibus test, pseudo-$R^2$ = 40.5%). The best model of *nosZ* contained fertilization and experiment duration as two non-important predictors (n = 36, $p = 0.006$ in omnibus test, pseudo-$R^2$ = 31.5%). The final urease model included biochar application rate as a single important moderator but it did not reduce the total non-sampling variance of the random-effects model (n = 64, $p < 0.001$ in omnibus test, pseudo-$R^2$ = 0%). Biochar application rate and experiment duration were important predictors in the final model of potential nitrification rate (n = 33, $p < 0.001$ in omnibus test, pseudo-$R^2$ = 63.0%). Feedstock (lignocellulose or wood) and fertilization were important predictors in the best model of cumulative $N_2O$ emissions (n = 63, $p <0.001$ in omnibus test, pseudo-$R^2$ = 59.4%).

**S2.4.2.3. P Cycling**

The final model of acid phosphatase activity contained three important predictors, pyrolysis temperature, soil pH, and experiment duration, as well as a non-important moderator, biochar application rate, but including these moderators did not reduce the total non-sampling variance of the random-effects model (n = 86, $p < 0.001$ in omnibus test, pseudo-$R^2$ = 0). The best alkaline phosphatase model included soil pH as a single, non-important predictor (n = 28, $p = 0.082$; pseudo-$R^2$ = 54.5%).

**S2.5. Publication Bias and Missing Data**

Overall, funnel plots suggest that publication bias existed for many microbial responses in the three-level random-effects models (Figure S3). Moderator analysis and model selection largely mitigated this issue, as reflected by the funnel plots and the results from Egger's test, Kendall's test, and Rosenthal's FSN test (Figure S4, Tables S9 and Table S10).

### S3. Supplementary Figures

**Figure S1.** Number of records per year retrieved from the Web of Science using "biochar soil amendment" as the search topic. Last accessed Jan 23, 2021.

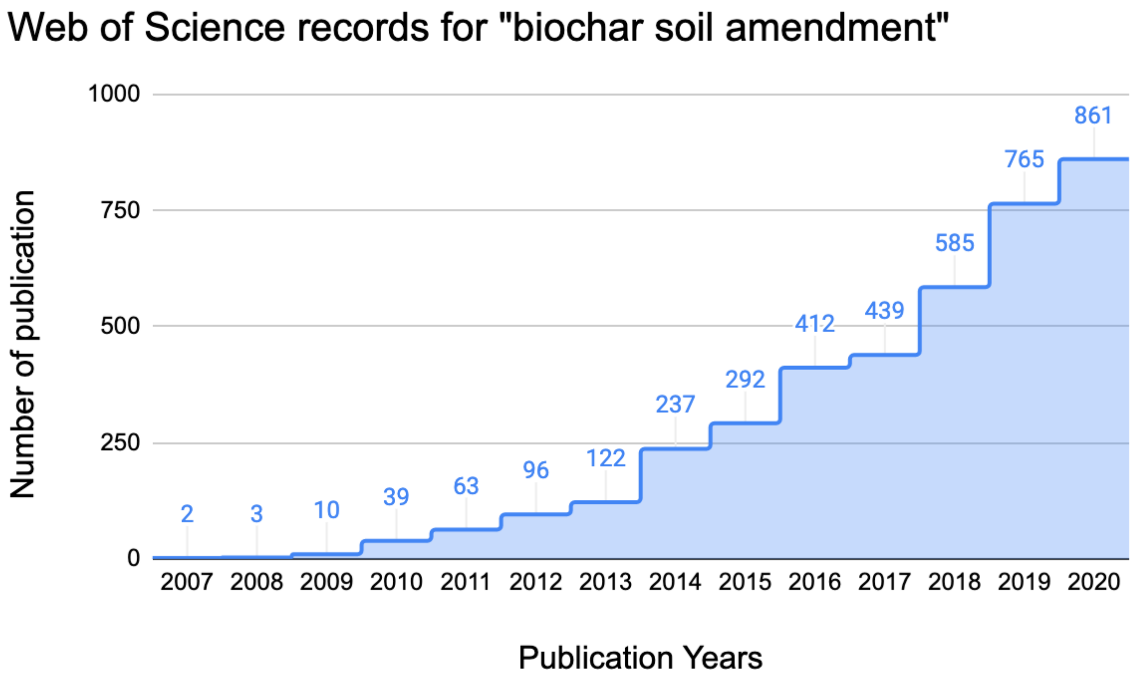

**Figure S2.** Significant single categorical moderators of soil microbiome responses. Mean effect sizes are presented as weighted LRR ± 95% confidence intervals. Effect sizes whose 95% confidence intervals overlapping with zero are considered as insignificant. Data presented here had <10 observations at certain moderator levels.

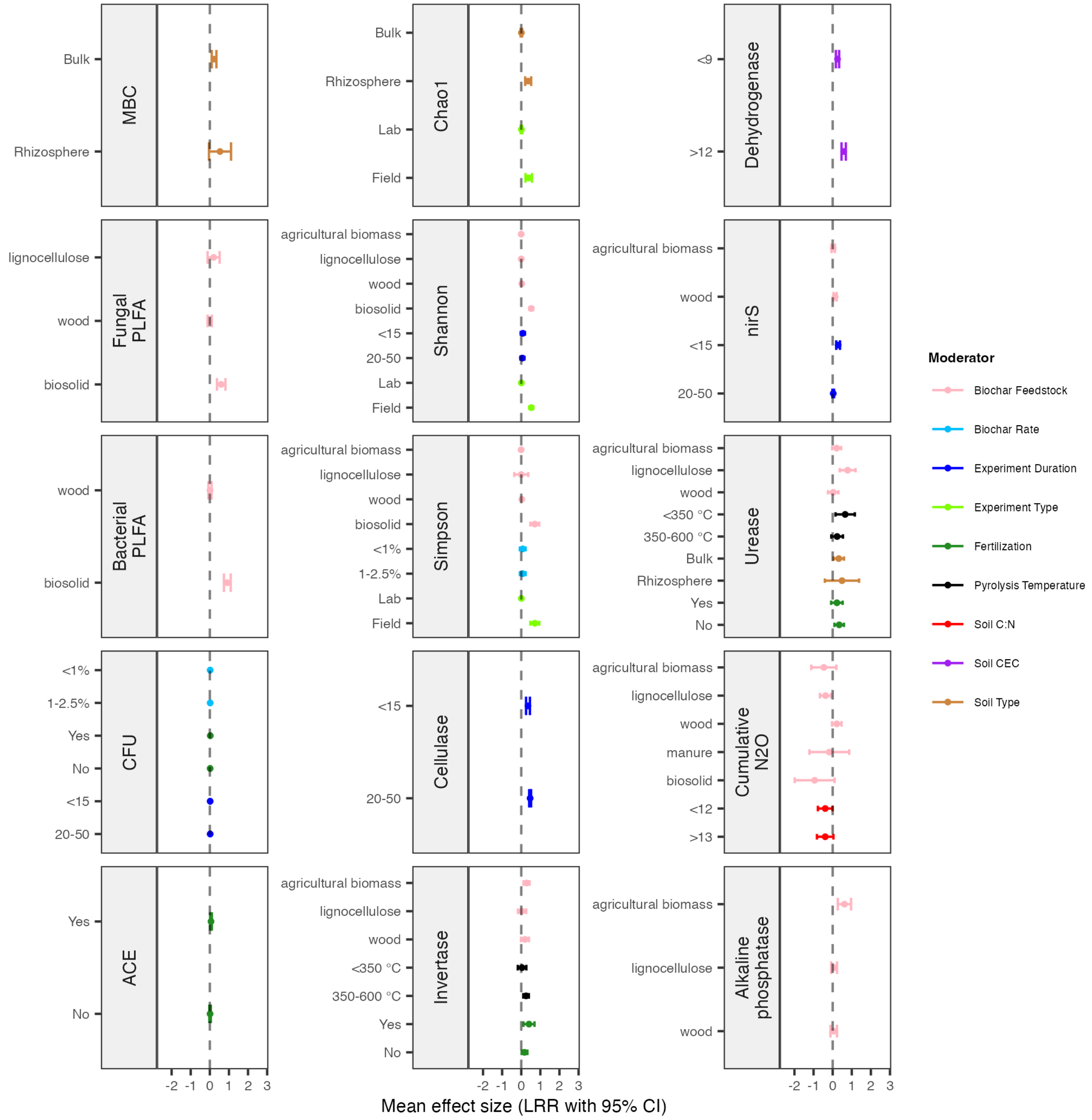

**Figure S3.** Funnel plots for global effect sizes of individual soil microbial responses before model selection. X-axis presents the effect size (LRR) and Y-axis shows the inverse standard error of the effect size as an index of precision. White and gray zones represent 90%, 95%, and 99% pseudo confidence interval regions.

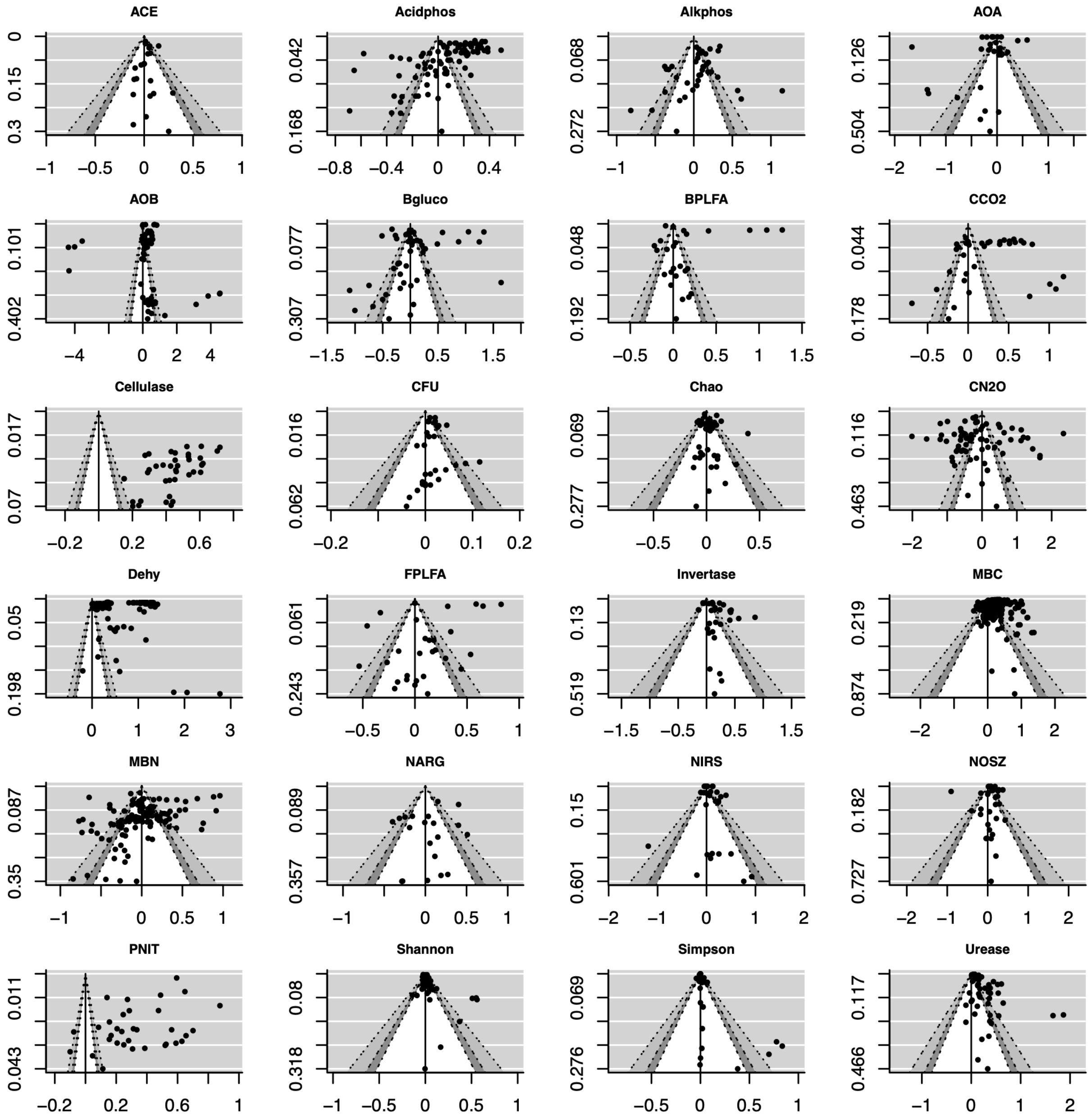

**Figure S4.** Funnel plots for global effect sizes of individual soil microbial responses in their final models. X-axis presents the effect size (LRR) and Y-axis shows the inverse standard error of the effect size as an index of precision. White and gray zones represent 90%, 95%, and 99% pseudo confidence interval regions.

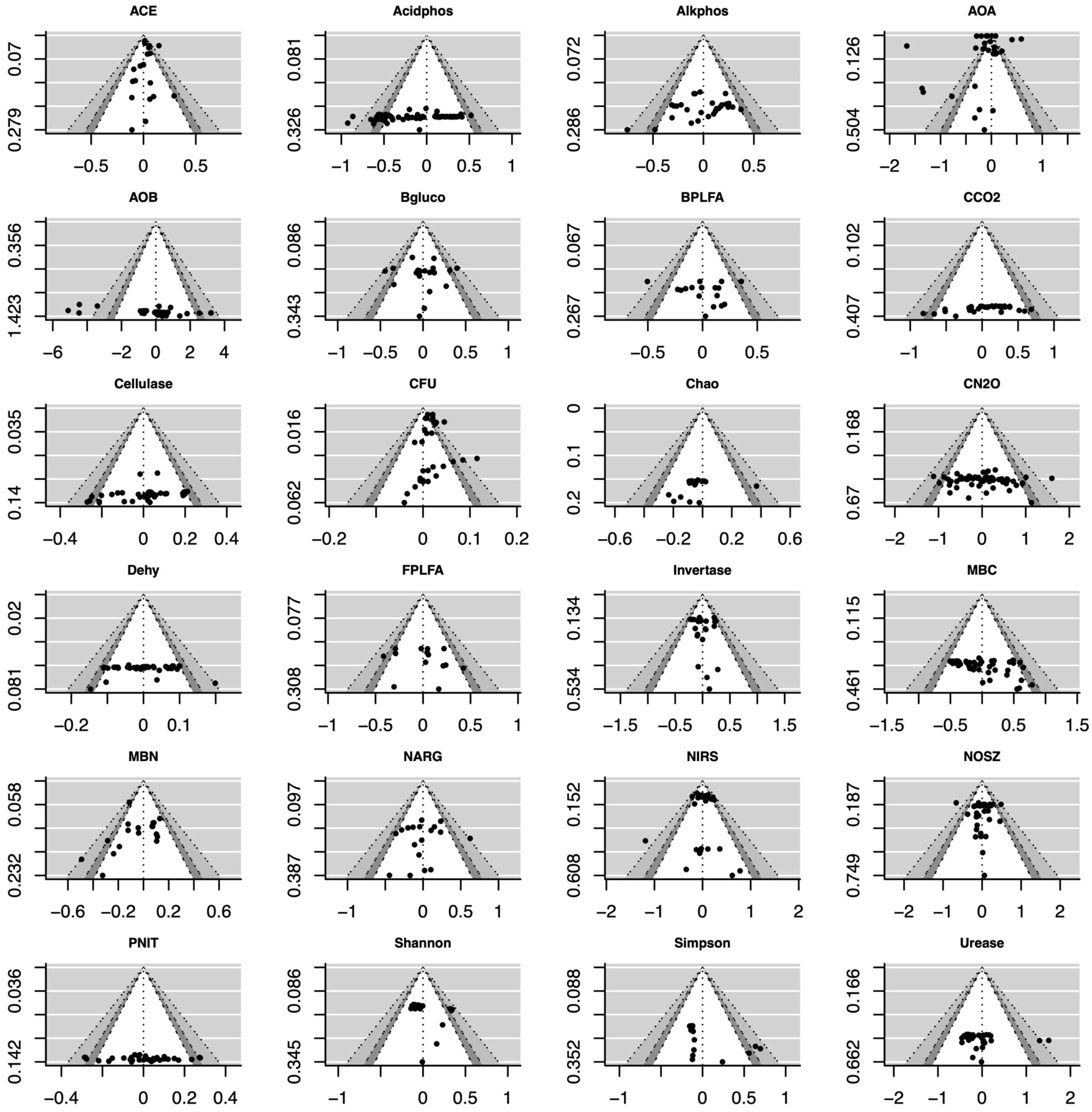

### S4. Supplementary Tables

**Table S1.** Keywords and keyword combinations used in the systematic literature search in the Web of Science database.

TS=(biochar AND microbiome)
TS=(biochar AND microbial communit*)
TS=(biochar AND (microb* OR microorg*))
TS=(biochar AND bacterial communit*)
TS=(biochar AND bacteri*)
TS=(biochar AND (bacterium OR bacteria OR bacterial))
TS=(biochar AND fungal communit*)
TS=(biochar AND fung*)
TS=(biochar AND (fungus OR fungi OR fungal))
TS=(biochar AND nutrient cycl* AND (microb* OR microorg*))
TS=(biochar AND nutrient metaboli* AND (microb* OR microorg*))
TS=(biochar AND carbon cycl* AND (microb* OR microorg*))
TS=(biochar AND carbon metaboli* AND (microb* OR microorg*))
TS=(biochar AND nitrogen cycl* AND (microb* OR microorg*))
TS=(biochar AND nitrogen metaboli* AND (microb* OR microorg*))
TS=(biochar AND phosphorus cycl* AND (microb* OR microorg*))
TS=(biochar AND phosphorus metaboli* AND (microb* OR microorg*))
TS=(biochar AND functional gene* AND (microb* OR microorg*))
TS=(biochar AND soil health AND (microb* OR microorg*))
TS=(biochar AND soil enzyme activity)
TS=(biochar AND soil enzyme)
TS=(biochar AND extracellular enzyme)
TS=(biochar AND exoenzyme)
TS=(biochar AND microbial ecology)
TS=(biochar AND community structure)
TS=(biochar AND community composition)
TS=(biochar AND community function)
TS=(biochar AND diversity AND (microb* OR microorg*))
TS=(biochar AND relative abundance AND (microb* OR microorg*))
TS=(biochar AND network AND (microb* OR microorg*))
TS=(biochar AND meta*omic*)

**Table S2.** A complete list of response variables obtained from the 61 primary studies. Variables with 20 or more observations were used in the meta-analysis.

| **Variable** | **Description** |
|---|---|
| ***Microbial abundance*** | |
| CFU | Number of colony forming unit (CFU) (counts/g soil) |
| MB | Microbial biomass carbon (mg/g soil) |
| MBC | Microbial biomass carbon (mg C/g soil) |
| MBN | Microbial biomass nitrogen (mg N/g soil) |
| Total PFLA | Phospholipid fatty acid (PLFA) abundance of all taxa (mg PLFA/g soil) |
| Bacteria | PFLA abundance of bacteria (mg PLFA/g soil) |
| Eubacteria | PFLA abundance of eubacteria (mg PLFA/g soil) |
| Anaerobe | PFLA abundance of anaerobes (mg PLFA/g soil) |
| Gram-positive bacteria | PFLA abundance of gram-positive bacteria (mg PLFA/g soil) |
| Gram-negative bacteria | PFLA abundance of gram-negative bacteria (mg PLFA/g soil) |
| Actinobactera | PFLA abundance of actinobacteria (mg PLFA/g soil) |
| Fungi | PFLA abundance of fungi (mg PLFA/g soil) |
| Eukaryote | PFLA abundance of eukaryotes (mg PLFA/g soil) |
| ***Microbial diversity*** | |
| OTU | Operational taxonomic unit (counts) |
| Observed species | Number of species (counts) |
| Shannon’s diversity | H’ = $-\sum_{i=1}^{n} p_i ln p_i$ ($p_i$ = relative abundance of species *i*) |
| Simpson’s diversity | Simpson's diversity index = $1-\sum_{i=1}^{k}\frac{{n_i}^2}{N}$ where N is total number of organisms, $n_i$ is the number of organisms of species *i*, and *k* is the number of species |
| ACE | ACE diversity index |
| Richness | Genetic richness based on number of unique amplicons identified (counts) |
| Evenness | Pielou's evenness index ($\frac{H'}{ln\ (Genetic\ Richness)}$) |
| Chao1 | Chao1 = number of species + (number of singletons)$^2$ / 2(number of doubletons) |
| ***Microbial community composition*** | |

| | |
|---|---|
| Acidobacteria | Phylum level abundance determined by 16s rRNA sequencing (%) |
| Actinobacteria | Phylum level abundance determined by 16s rRNA sequencing (%) |
| Armatimonadetes | Phylum level abundance determined by 16s rRNA sequencing (%) |
| Bacteroidetes | Phylum level abundance determined by 16s rRNA sequencing (%) |
| Chlamydiae | Phylum level abundance determined by 16s rRNA sequencing (%) |
| Chlorobi | Phylum level abundance determined by 16s rRNA sequencing (%) |
| Chloroflexi | Phylum level abundance determined by 16s rRNA sequencing (%) |
| Cyanobacteria | Phylum level abundance determined by 16s rRNA sequencing (%) |
| Deinococcus-Thermus | Phylum level abundance determined by 16s rRNA sequencing (%) |
| Elusimicrobia | Phylum level abundance determined by 16s rRNA sequencing (%) |
| Firmicutes | Phylum level abundance determined by 16s rRNA sequencing (%) |
| Gemmatimonodetes | Phylum level abundance determined by 16s rRNA sequencing (%) |
| Latescibacteria | Phylum level abundance determined by 16s rRNA sequencing (%) |
| Nitrospira | Phylum level abundance determined by 16s rRNA sequencing (%) |
| Parcubacteria | Phylum level abundance determined by 16s rRNA sequencing (%) |
| Planctomycetes | Phylum level abundance determined by 16s rRNA sequencing (%) |
| Proteobacteria | Phylum level abundance determined by 16s rRNA sequencing (%) |
| Saccharibacteria | Phylum level abundance determined by 16s rRNA sequencing (%) |
| Spirochaetes | Phylum level abundance determined by 16s rRNA sequencing (%) |
| Synergistetes | Phylum level abundance determined by 16s rRNA sequencing (%) |
| Tenericutes | Phylum level abundance determined by 16s rRNA sequencing (%) |
| Thaumarchaeota | Phylum level abundance determined by 16s rRNA sequencing (%) |
| Thermotogae | Phylum level abundance determined by 16s rRNA sequencing (%) |
| Verrucomicrobia | Phylum level abundance determined by 16s rRNA sequencing (%) |
| BRC1 | Phylum level abundance determined by 16s rRNA sequencing (%) |
| Microgenomates | Phylum level abundance determined by 16s rRNA sequencing (%) |
| OD1 | Phylum level abundance determined by 16s rRNA sequencing (%) |
| OP11 | Phylum level abundance determined by 16s rRNA sequencing (%) |
| TM7 | Phylum level abundance determined by 16s rRNA sequencing (%) |
| WD272 | Phylum level abundance determined by 16s rRNA sequencing (%) |

| | |
|---|---|
| WS3 | Phylum level abundance determined by 16s rRNA sequencing (%) |
| Others | Phylum level abundance determined by 16s rRNA sequencing (%) |
| Unclassified | Phylum level abundance determined by 16s rRNA sequencing (%) |
| ***Microbial community function*** | |
| Carbon cycling | |
| Enzymes | |
| β-glucosidase | Activity of β-glucosidases that catalyze the hydrolysis of the glycosidic bonds in β-D-glucosides (μg *p*-nitrophenol (PNP)/g soil/h) |
| Cellulase | Activity of cellulases that catalyze the decomposition of cellulose and some related polysaccharides (μg glucose/g soil/h) |
| Invertase | Activity of invertases that catalyze the hydrolysis of sucrose into fructose and glucose (μg glucose/g soil/h) |
| Nitrogen cycling | |
| Genes | |
| *narG* | Copy number of nitrate reductase gene *narG* (copies/g soil) |
| *nirK* | Copy number of nitrite reductase gene *nirK* (copies/g soil) |
| *nirS* | Copy number of nitrite reductase gene *nirS* (copies/g soil) |
| *nosZ* | Copy number of nitrous oxide reductase gene *nosZ* (copies/g soil) |
| Enzymes | |
| Protease | Activity of proteases that catalyze the hydrolysis of proteins (μg tyrosine/g soil/h or μg $NH4^+$-N/g soil/h) |
| Urease | Activity of ureases that catalyze the hydrolysis of urea (μg $NH4^+$-N/g soil/h) |
| Population | |
| AOA | Number of ammonia-oxidizing archaea, mostly quantified by measuring the single copy gene *amoA* (counts/g soil) |
| AOB | Number of ammonia-oxidizing bacteria, mostly quantified by measuring the single copy gene *amoA* (counts/g soil) |
| Phosphorus cycling | |
| Enzymes | |
| Phosphomonoesterase | Activity of phosphomonoesterases that catalyze the hydrolysis of phosphoric monoesters to phosphate (μg *p*-nitrophenol (PNP)/g soil/h) |
| Acid phosphatase | Activity of acid phosphatases that catalyze the hydrolysis of both esters and anhydrides of phosphoric acid (μg *p*-nitrophenol (PNP)/g soil/h) |

| | |
|---|---|
| Alkaline phosphatase | Activity of alkaline phosphatases that catalyze the hydrolysis of both esters and anhydrides of phosphoric acid (µg *p*-nitrophenol (PNP)/g soil/h) |
| Oxidation-reduction | |
| Enzymes[a] | |
| Catalase | Activity of enzyme catalases that catalyze the conversion of hydrogen peroxide to water and oxygen (µg $H_2O_2$/g soil/h) |
| Dehydrogenase | Activity of dehydrogenases that catalyze the transfer of hydrogen and electrons between organic compounds (µg iodonitrotetrazolium formazan (INTF)/g soil/h) |
| ***Ecosystem processes*** | |
| AWCD | Average well color development, an index of substrate utilization |
| FDA | Fluorescein diacetate hydrolysis by a variety of enzymes including esterases, lipases, and proteases, an indicator of organic matter turnover by total microbial activity (µg fluorescein/g soil/h) |
| Metabolic quotient ($qCO_2$) | Respiration-to-biomass ratio, an index of ecosystem development (during which it supposedly declines) and disturbance (due to which it supposedly increases) that is conceptually based on Odum's theory of ecosystem succession (µg $CO_2$-C/mg microbial C/h) |
| Potential nitrification rate | Overall nitrification rate measured based on (nmol N/g dry soil/h) |
| Soil $CO_2$ emissions | Soil $CO_2$ emissions (mg/g soil) |
| Soil cumulative $CO_2$ | Cumulative $CO_2$ emissions over a given time period (mg/g soil) |
| Soil $CO_2$ equivalents | Greenhouse gas emissions expressed in $CO_2$ of $CO_2$ , measure of how much a gas contributes to climate change, relative to CO2 (Kg/ha/yr) |
| Soil $CH_4$ emissions | Soil $CH_4$ emissions (mg/g soil) |
| Soil cumulative $CH_4$ | Cumulative $CH_4$ emissions over a given time period (mg/g soil) |
| Soil $N_2O$ emissions | Soil $N_2O$ emissions (mg/g soil) |
| Soil cumulative $N_2O$ | Cumulative $N_2O$ emissions over a given time period (mg/g soil) |
| ***Plant physiology*** | |
| Height | Plant Height (cm) |
| Fresh weight | Fresh weight of plant biomass (g/plant) |
| Dry weight | Dry weight of plant biomass (g/plant) |
| Seed yield | Number of seeds yielded per plant |
| Shoot dry biomass | Dry weight of plant shoot biomass (g/plant) |
| Root dry biomass | Dry weight of plant root biomass (g/plant) |

| | |
|---|---|
| Root colonization by arbuscular mycorrhizal fungi (AMF) | Mycorrhizal root colonization measured visually with microscope (% root length colonized) |
| ***Soil property*** | |
| pH | Soil pH |
| Moisture content | Volumetric water content of soil (%) |
| Electrical conductivity (EC) | Measure of salinity of soil (mS/cm) |
| Total carbon (TC) | Total soil carbon (%) |
| Organic carbon (OC) | Total soil organic carbon (%) |
| Total nitrogen (TN) | Total soil nitrogen (%) |
| Ammonium ($NH_4^+$) | Amount of ammonium in soil (mg/kg) |
| Nitrate ($NO_3^-$) | Amount of nitrate in soil (mg/kg) |
| Carbon nitrogen ratio (C:N) | Ratio of total C to total N in soil |
| Total phosphorus (TP) | Total soil phosphorus (%) |
| Available phosphorus (P) | Amount of inorganic phosphorus available for plant uptake (mg/kg) |
| Available potassium (K) | Amount of potassium available for plant uptake (mg/kg) |

[a]Oxidation-reduction enzymes are merged with C-cycling enzymes in plotting.

**Table S3.** A complete list of predictor variables (moderators) used in this study. To facilitate cross-study comparisons, moderators were assigned into categorical groups. In the later on model selection, soil pH, soil C:N, experiment duration, and biochar application rate were included as continuous predictors. This decision was made based on the continuous nature of these parameters.

| Variable | Abbreviation | Units | Levels | Description |
|---|---|---|---|---|
| ***Soil property*** | | | | |
| Texture | TEXT | USDA classification | | |
| pH | PH | N/A | A<br>B<br>C | <5<br>5-8 (inclusive)<br>>8 |
| Cation exchange capacity | CEC | cmol/kg | A<br>B | <9<br>>12 |
| C:N ratio | CN | N/A | A<br>B | <12<br>>13 |
| Total C | TC | % | A<br>B<br>C | <1.6<br>1.6-4.0 (inclusive)<br>>9 |
| Soil organic C | SOC | % | A<br>B | <0.4<br>>0.5 |
| Soil organic matter | SOM | % | A<br>B | <1.8<br>>2 |
| Total N | TN | % | A<br>B<br>C | <0.1<br>0.1-0.5 (inclusive)<br>≥0.8 |
| $NH_4$-N | NH4 | mg/kg | A<br>B | <10<br>>14 |
| $NO_3$-N | NO3 | mg/kg | A<br>B | <10<br>>14 |
| Total P | TP | % | A<br>B | <0.08<br>>0.1 |
| Available P | AP | mg/kg | A<br>B<br>C | <9<br>9-14 (inclusive)<br>>15 |
| Available K | AK | mg/kg | A<br>B | <70<br>>71 |
| ***Biochar property*** | | | | |
| Feedstock | FEED | N/A | | |
| Production technique | Type | N/A | | |
| Production temperature | TEMP | °C | low<br>med | ≤350<br>350-600 |

| | | | | |
|---|---|---|---|---|
| | | | high | ≥600 |
| pH | B_PH | N/A | A<br>B<br>C | <7<br>7-9.8 (inclusive)<br>>9.8 |
| Ash content | B_ASH | % | A<br>B<br>C | <6<br>6-20 (inclusive)<br>>20 |
| C:N ratio | B_CN | N/A | A<br>B<br>C | <30<br>30-60 (inclusive)<br>>70 |
| C content | B_TC | % | A<br>B<br>C | <33<br>34-59 (inclusive)<br>≥60 |
| Organic C | B_OC | mg/kg | A<br>B | <25<br>>30 |
| N content | B_TN | % | A<br>B<br>C | <0.4<br>0.4-1.0 (inclusive)<br>>1.1 |
| NO3-N | B_NO3 | mg/kg | A<br>B | <5<br>>6 |
| NH4-N | B_NH4 | mg/kg | A<br>B | <11<br>>14 |
| P content | B_TP | % | A<br>B<br>C | <0.1<br>0.1-0.5 (including 0.1)<br>≥0.5 |
| Available P | B_AP | mg/kg | A<br>B | <40<br>>200 |
| Available K | B_AK | mg/kg | A<br>B | <27<br>>160 |
| ***Experiment design*** | | | | |
| Experiment type | EXP | N/A | Laboratory<br>Field | |
| Biochar application rate | RATE | % | low<br>med<br>high | <1<br>1-2.5 (inclusive)<br>>2.5 |
| Fertilizer addition | FERT | N/A | Yes<br>No | |
| Duration | DAY | Days | A<br>B<br>C<br>D | ≤15<br>20-50 (inclusive)<br>60-90 (inclusive)<br>≥100 |
| Soil location | SOIL | N/A | Rhizosphere<br>Bulk | |
| Incubation temperature | INCUBATION | ° C | A<br>B | <25<br>≥25 |
| Maintained water content | WHC | %WHC[a] | A<br>B<br>C | ≤60<br>70-80<br>100 |

[a]WHC, water holding capacity.

**Table S4.** Raw data extracted from the 61 primary studies. Predictor variables are assigned into categorical groups as detailed in Table S3. In the later on model selection, soil pH, soil C:N, experiment duration, and biochar application rate were included as continuous predictors (see excel sheet).

**Table S5.** Full results of three-level random-effects models for the 24 microbial responses with at least 20 observations. *n*, number of observations; CI, confident interval; level-2 or level-3 $I^2$, the proportion of variation in effect sizes explained by level-2 or level-3 variance. Significant p-values ($\alpha$ = 0.05) are in bold. Also see excel sheet Table S5.

| **Soil microbiome parameter** | ***n*** | **Global estimate (LRR)** | **CI** | ***p*-value** | **Cochran's Q** | ***Q p*-value** | **Level-2 $I^2$ (%)** | **Level-3 $I^2$ (%)** |
|---|---|---|---|---|---|---|---|---|
| **Abundance** | | | | | | | | |
| MBC | 202 | 0.235 | 0.120 | **<0.001** | 11360.385 | **<0.001** | 29.044 | 70.423 |
| MBN | 116 | -0.041 | 0.164 | 0.625 | 2499.706 | **<0.001** | 52.405 | 43.008 |
| Fungal PLFA | 30 | 0.158 | 0.246 | 0.198 | 4221.791 | **<0.001** | 43.161 | 55.909 |
| Bacterial PLFA | 27 | 0.246 | 0.460 | 0.281 | 22188.952 | **<0.001** | 13.395 | 86.505 |
| CFU | 32 | 0.017 | 0.006 | **<0.001** | 46.941 | **0.033** | 35.700 | 0.000 |
| **Diversity** | | | | | | | | |
| ACE | 23 | 0.029 | 0.040 | 0.143 | 26.944 | 0.213 | 33.884 | 7.655 |
| Chao1 | 47 | 0.049 | 0.082 | 0.235 | 189.106 | **<0.001** | 7.944 | 84.570 |
| Shannon | 60 | 0.053 | 0.095 | 0.266 | 357.069 | **<0.001** | 0.667 | 98.990 |
| Simpson | 38 | 0.080 | 0.150 | 0.286 | 265.694 | **<0.001** | 0.005 | 99.994 |
| **Function and process** | | | | | | | | |
| $\beta$-Glucosidase | 48 | 0.002 | 0.355 | 0.991 | 5708.195 | **<0.001** | 25.460 | 73.792 |

| | | | | | | | | |
|---|---|---|---|---|---|---|---|---|
| Cellulase | 40 | 0.442 | 0.047 | **<0.001** | 525.078 | **<0.001** | 92.608 | 0.000 |
| Invertase | 31 | 0.192 | 0.140 | **0.009** | 414.543 | **<0.001** | 49.839 | 46.448 |
| Dehydrogenase | 128 | 0.610 | 0.365 | **0.001** | 236695.207 | **<0.001** | 53.396 | 46.569 |
| Cumulative $CO_2$ | 38 | 0.202 | 0.380 | 0.288 | 1940.547 | **<0.001** | 13.440 | 85.962 |
| AOA | 28 | -0.192 | 0.287 | 0.182 | 9589.758 | **<0.001** | 84.022 | 15.953 |
| AOB | 61 | 0.357 | 1.151 | 0.537 | 47583.407 | **<0.001** | 49.344 | 50.649 |
| *narG* | 20 | 0.022 | 0.279 | 0.871 | 76.690 | **<0.001** | 41.239 | 36.461 |
| *nirS* | 28 | 0.079 | 0.072 | **0.032** | 987878.576 | **<0.001** | 95.162 | 4.837 |
| *nosZ* | 36 | 0.071 | 0.115 | 0.216 | 8111.059 | **<0.001** | 91.131 | 8.838 |
| Urease | 74 | 0.332 | 0.248 | **0.009** | 1331.317 | **<0.001** | 1.977 | 97.990 |
| Cumulative $N_2O$ | 71 | -0.136 | 0.288 | 0.350 | 3156.402 | **<0.001** | 79.622 | 18.910 |
| Potential Nitrification Rate | 33 | 0.342 | 0.083 | **<0.001** | 6923.759 | **<0.001** | 99.608 | 0.000 |
| Acid Phosphatase | 87 | 0.033 | 0.110 | 0.559 | 4334.422 | **<0.001** | 45.500 | 53.282 |
| Alkaline Phosphatase | 38 | 0.173 | 0.196 | 0.082 | 324.834 | **<0.001** | 51.160 | 43.738 |

**Table S6.** Full summary of statistics of all the three-level mixed-effects models including one single categorical moderator (see excel sheet). For each microbial response variable, moderators were included when data were available. Occasionally, the number of observations at a particular moderator level dropped below 10. $R^2$ reflects the proportional reduction in the total heterogeneity after including the particular moderator in the model; negative $R^2$ values were truncated to zero.

**Table S7.** Importance of individual predictors to soil microbial responses (see excel sheet). Data presented here include model-averaged parameter estimates, which are weighted averages of the model coefficients across the various models (with weights equal to the model probabilities), unconditional variance of the model-averaged estimates, importance, and the confidence intervals built under alpha risk of 0.05.

**Table S8.** Full summary of statistics of the final three-level mixed-effects models (see excel sheet). Four predictors (soil pH, soil C:N, experiment duration, biochar application rate) were included as continuous moderators. Multicollinearity of candidate moderators was evaluated using variance inflation factor (VIF). The best model was selected based on corrected Akaike information criterion (AICc). $R^2$ reflects the proportional reduction in the total variance after including a particular moderator in the random-effects model; for negative $R^2$ values, they were truncated to zero.

**Table S9.** Results of Egger's linear regression test, Kendall's tau rank correlation test, and Rosenthal's fail-safe (FSN) test for assessing publication bias in the three-level random-effects models including one single categorical moderator (see excel sheet).

**Table S10.** Results of Egger's linear regression test, Kendall's tau rank correlation test, and Rosenthal's fail-safe (FSN) test for assessing publication bias in the final best three-level random-effects models including multiple moderators (see excel sheet).
